# Manipulating the EphB4-ephrinB2 axis to reduce metastasis in HNSCC

**DOI:** 10.1101/2024.07.21.604518

**Authors:** Khalid N.M. Abdelazeem, Diemmy Nguyen, Sophia Corbo, Laurel B. Darragh, Mike W. Matsumoto, Benjamin Van Court, Brooke Neupert, Justin Yu, Nicholas A. Olimpo, Douglas Grant Osborne, Jacob Gadwa, Richard B. Ross, Alexander Nguyen, Shilpa Bhatia, Mohit Kapoor, Rachel S. Friedman, Jordan Jacobelli, Anthony J. Saviola, Michael W. Knitz, Elena B. Pasquale, Sana D. Karam

**Affiliations:** Department of Radiation Oncology, University of Colorado Denver, Anschutz Medical Campus, Aurora, CO, USA; Radiation Biology Research Department, National Center for Radiation Research and Technology, Egyptian Atomic Energy Authority, Cairo, Egypt; Department of Immunology and Microbiology, University of Colorado Anschutz Medical Campus, Aurora, CO, USA; Cancer Center, Sanford Burnham Prebys Medical Discovery Institute, La Jolla, CA, USA; Department of Otolaryngology - Head and Neck Surgery, University of Colorado Anschutz Medical Campus, Aurora, CO, USA; Department of Dermatology, University of Colorado Anschutz Medical Campus, Aurora, CO, USA; Krembil Research Institute, University Health Network, and University of Toronto, Toronto, Ontario, Canada; Barbara Davis Research Center, University of Colorado Anschutz Medical Campus, Aurora, CO, USA; Department of Biochemistry and Molecular Genetics, University of Colorado Anschutz Medical Center, Aurora, CO, USA

## Abstract

The EphB4-ephrinB2 signaling axis has been heavily implicated in metastasis across numerous cancer types. Our emerging understanding of the dichotomous roles that EphB4 and ephrinB2 play in head and neck squamous cell carcinoma (HNSCC) poses a significant challenge to rational drug design. We find that EphB4 knockdown in cancer cells enhances metastasis in preclinical HNSCC models by augmenting immunosuppressive cells like T regulatory cells (Tregs) within the tumor microenvironment. EphB4 inhibition in cancer cells also amplifies their ability to metastasize through increased expression of genes associated with epithelial mesenchymal transition and hallmark pathways of metastasis. In contrast, vascular ephrinB2 knockout coupled with radiation therapy (RT) enhances anti-tumor immunity, reduces Treg accumulation into the tumor, and decreases metastasis. Notably, targeting the EphB4-ephrinB2 signaling axis with the engineered EphB4 ligands EFNB2-Fc-His and Fc-TNYL-RAW-GS reduces local tumor growth and distant metastasis in a preclinical model of HNSCC. Our data suggest that targeted inhibition of vascular ephrinB2 while avoiding inhibition of EphB4 in cancer cells could be a promising strategy to mitigate HNSCC metastasis.

## Introduction

Head and neck squamous cell carcinoma (HNSCC) is a formidable challenge in oncology and stands as the sixth most prevalent cancer type globally with a 5-year survival rate below 50% (Duprez et al., 2017; Fuereder, 2022; Sun et al., 2022). Despite advances in treatment modalities, the prognosis remains notably poor, particularly for patients with advanced disease. Approximately 10% of HNSCC patients present with distant metastases, and up to 30% additional patients develop distant metastases as their disease progresses (Duprez et al., 2017). Patients with distant metastases have a median overall survival of only 10 months. These grim statistics underscore the urgent need for innovative therapeutic strategies specifically targeting distant metastases in the context of HNSCC (Duprez et al., 2017).

Metastasis is a complex process through which cancer cells escape from the primary tumor and seed distant sites (Pisani et al., 2020). The process of metastasis involves a series of sequential steps, including invasion, intravasation into blood or lymphatic vessels, circulation, extravasation at distant sites, and colonization, each of which is influenced by complex interactions between tumor cells, the vascular network, and the immune microenvironment (Fares et al., 2020). The tumor vasculature is a physical barrier that can regulate infiltration of cancer and immune cells into the tumor, making it an attractive target to inhibit metastasis (Nagl et al., 2020). Recent evidence indicates that adhesion and co-stimulatory molecules expressed by the tumor endothelium can promote cancer cell and lymphocyte adhesion, transmigration, or survival, thereby modulating the local and systemic anti-tumor response (Fu et al., 2014; Motz et al., 2014).

Numerous genes have been implicated in metastasis, including the Eph receptor and ephrin gene families (Pasquale, 2010). Eph receptors form the largest family of receptor tyrosine kinases (RTKs); they bind to membrane-bound ephrin ligands, and both are capable of signal transduction (Pfaff et al., 2008). Eph receptors and ephrins play crucial roles during embryogenesis, particularly in processes such as axon guidance, cell migration, angiogenesis, and hematopoiesis (Pasquale, 2010). In adulthood, Eph receptors and ephrins are expressed during tissue repair, wound healing, and pathologies such as immune diseases and cancer, where they function to mediate neo-angiogenesis, fibrosis, inflammation, and immunosuppression (Funk & Orr, 2013; Lu et al., 2017; Wu & Luo, 2005).

EphB4 and ephrinB2 are a receptor-ligand pair that has been heavily studied for their role in metastasis across numerous cancer types (Alam et al., 2008; Broggini et al., 2020; Héroult et al., 2010; Li et al., 2019). We have reported that within the HNSCC tumor microenvironment (TME), EphB4 is predominantly expressed in cancer cells and its knockdown or knockout drives local progression by increasing immunosuppressive immune cells in the TME (Bhatia et al., 2022). In contrast, ephrinB2 is highly expressed on the vascular endothelium, and its knockout in this compartment suppresses tumor growth, decreases intratumoral infiltration of regulatory T cells (Tregs), and increases activation of CD8+ T cells (Bhatia et al., 2022). These data suggest that cancer cell EphB4 and vascular ephrinB2 play dichotomous roles in HNSCC progression. However, our previous studies were limited to examining the role of EphB4-ephrinB2 signaling in mediating local tumor growth. Given the prevalence of distant metastases in HNSCC and the limited translational therapeutics in that space, we sought to investigate how this signaling axis impacts metastatic spread.

Ongoing research efforts aim at investigating the impact of manipulating EphB4 and ephrinB2 signaling pathways in metastasis and tumor progression. Our findings reveal that EphB4 knockdown in cancer cells leads to an increase in the development of distant metastases, characterized by a pro-metastatic cancer cell phenotype, systemic immunosuppression, and enhanced infiltration of CD4+ T cells and Tregs into the tumor. We also show that ephrinB2 knockout in the vascular endothelium decreases local tumor growth and metastasis. This effect is mediated by improved effector T cell activity and an increase in T cell expansion in the tumor draining lymph nodes. Finally, we test two novel agents, ephrinB2-Fc and Fc-TNYL-RAW-GS, for their ability to overcome distant metastases. Overall, our study underscores the significance of EphB4 and ephrinB2 signaling pathways in HNSCC metastasis and highlights the therapeutic potential of targeting this axis to disrupt metastasis in HPV-unrelated HNSCC.

## Results

### Knockdown of EphB4 in HNSCC cancer cells promotes distant metastasis

We had previously demonstrated that EphB4 is predominantly expressed in cancer cells in various HPV-negative models of HNSCC and that its knockdown or knockout accelerates local tumor growth in the absence of radiation therapy (RT) (Bhatia et al., 2022). Here, we sought to determine the effects of EphB4 on distant metastasis using two orthotopic models of HNSCC in the context of RT: the MOC2 (C57BL/6J) cell line and the LY2 (BALB/c) cell line, with either control or EphB4 shRNA knockdown (Bhatia et al., 2022). Concordant with our previous findings (Bhatia et al., 2022), downregulation of EphB4 in cancer cells increases local tumor growth in both models (Suppl. Figure 1A-F). For metastasis studies, MOC2 cells were implanted in the buccal mucosa while LY2 cells were implanted in the floor of the mouth. Tumors were irradiated and imaged with computed tomography (CT) scans to monitor disease progression (Figure 1A, D, Suppl. Figure 1G). Interestingly, EphB4 knockdown in cancer cells was associated with an increase in the incidence of lung (MOC2, Figure 1B, C) or mediastinal (LY2, Figure 1E, F) metastases. Histological validation of the metastatic lesions was performed based on hematoxylin and eosin (H&E) staining of lung tissues (Suppl. Figure 1H). These findings indicate that EphB4 plays a multifaceted role in tumor biology, and that its downregulation in cancer cells increases not only local progression, but also distant spread.

**Figure 1:**
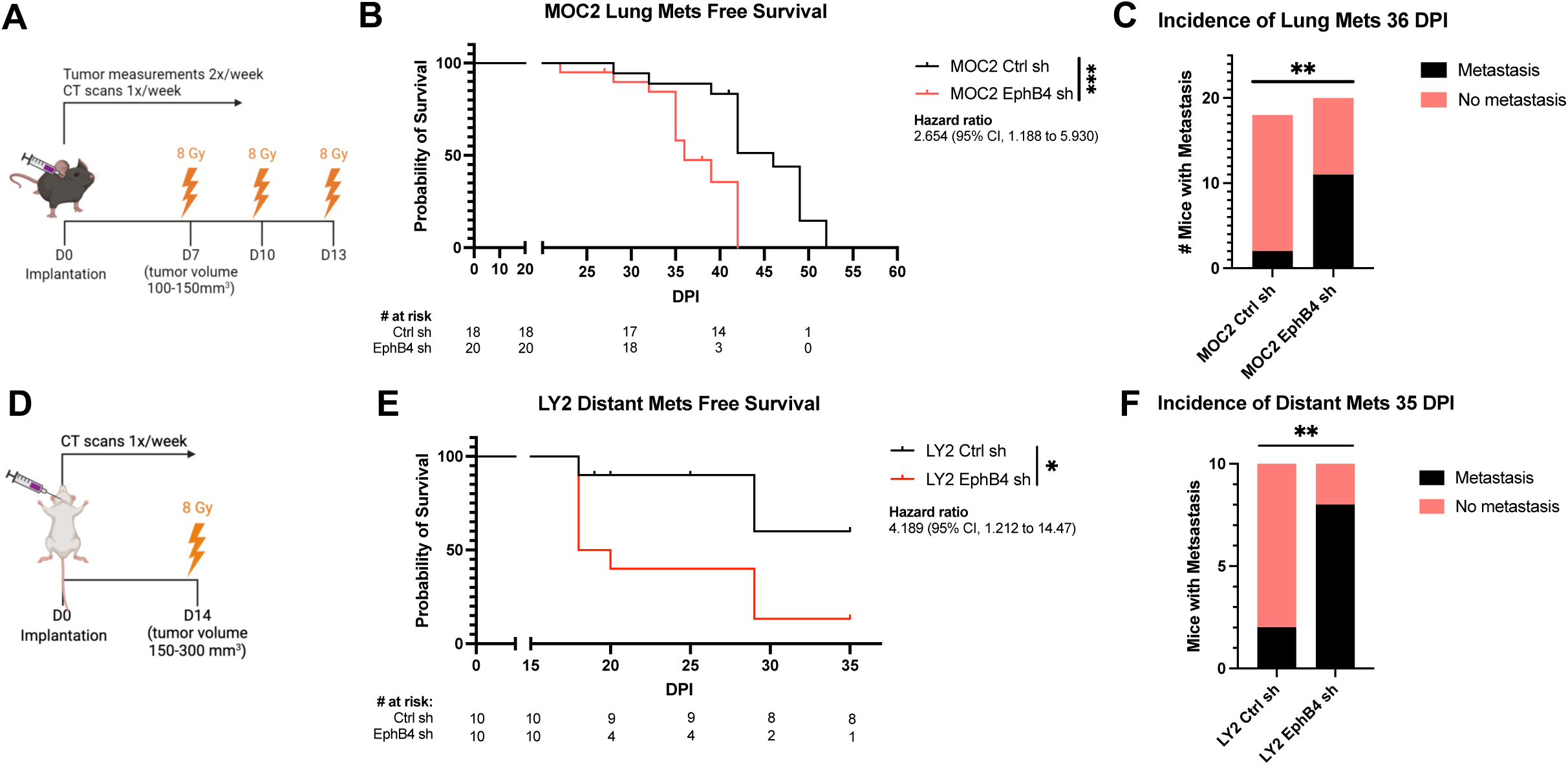
Knockdown of EphB4 in HNSCC cancer cells promotes distant metastasis. (A) Schematic showing experimental design for C57BL/6J mice implanted with 100k MOC2 cancer cells. 3 fractions of 8 Gray (Gy) radiation therapy (RT) were given as indicated. (B) Kaplan-Meier curves showing lung metastasis free survival of MOC2 control (Ctrl) shRNA (sh) versus EphB4 shRNA tumors implanted in C57BL/6J mice. Numbers at risk indicate mice that were alive without metastases at specified timepoints. (C) Contingency table quantifying the incidence of lung metastasis detected by computed tomography (CT) scans in C57BL/6 mice implanted with MOC2 cancer cells by 36 days post-implantation (DPI). (D) Schematic showing experimental design for BALB/c mice implanted with 100k LY2 cancer cells. 3 fractions of 8 Gy RT were given as indicated. (E) Kaplan-Meier curves showing distant metastasis free survival of LY2 Ctrl shRNA vs LY2 EphB4 shRNA tumors implanted in BALB/c mice. Numbers at risk indicate mice that were alive without metastases at specified timepoints. (F) Contingency table quantifying the incidence of distant metastases detected by CT scans in BALB/c mice implanted with LY2 Ctrl shRNA or EphB4 shRNA tumors by 37 DPI. For Kaplan-Meier survival curves, significance was determined by a log-rank Mantel-Cox test. For contingency tables indicating the incidence of metastases, significance was determined by a Chi-square test. Significance was determined if the *p*-value was <0.05*, <0.01**, <0.001***, and <0.0001****. *p*-values are indicated for the figures B \*\*\**p* = 0.0009, C \*\**p* = 0.0044, E \**p* = 0.0200, F \*\**p* = 0.0073.

### Loss of EphB4 in cancer cells induces protein dysregulation concomitant with increased metastatic capacity

To investigate whether EphB4 inhibition enhances metastasis through a canonical cancer-cell centered mechanism, we explored intrinsic changes induced by loss of EphB4 in the cancer cell using RNA sequencing. Comparison of transcriptomic changes between MOC2 control and EphB4 knockout cell lines showed higher expression of TGF beta receptors 1 and 2, NOTCH1, and FOXC1, a transcription factor implicated in cancer cell plasticity, treatment resistance, invasion, and epithelial-mesenchymal transition (EMT) (Figure 2A) (Ray et al., 2021; Zhu et al., 2017). We also observed an increase in metastasis-related hallmark pathways including TNF-alpha signaling via NF-κB, MYC, IL6/JAK/STAT3 signaling, and KRAS in MOC2 EphB4 knockdown cells (Figure 2B) (Boutin et al., 2017; Johnson et al., 2018; Maddipati et al., 2022; Schulze et al., 2020; Tang et al., 2017). To further investigate the effect of EphB4 on cancer cell migration, we employed Transwell Boyden chamber assays. We observed that EphB4 knockdown cancer cells had increased migratory capacity compared to control cells *in vitro* (Figure 2C).

**Figure 2:**
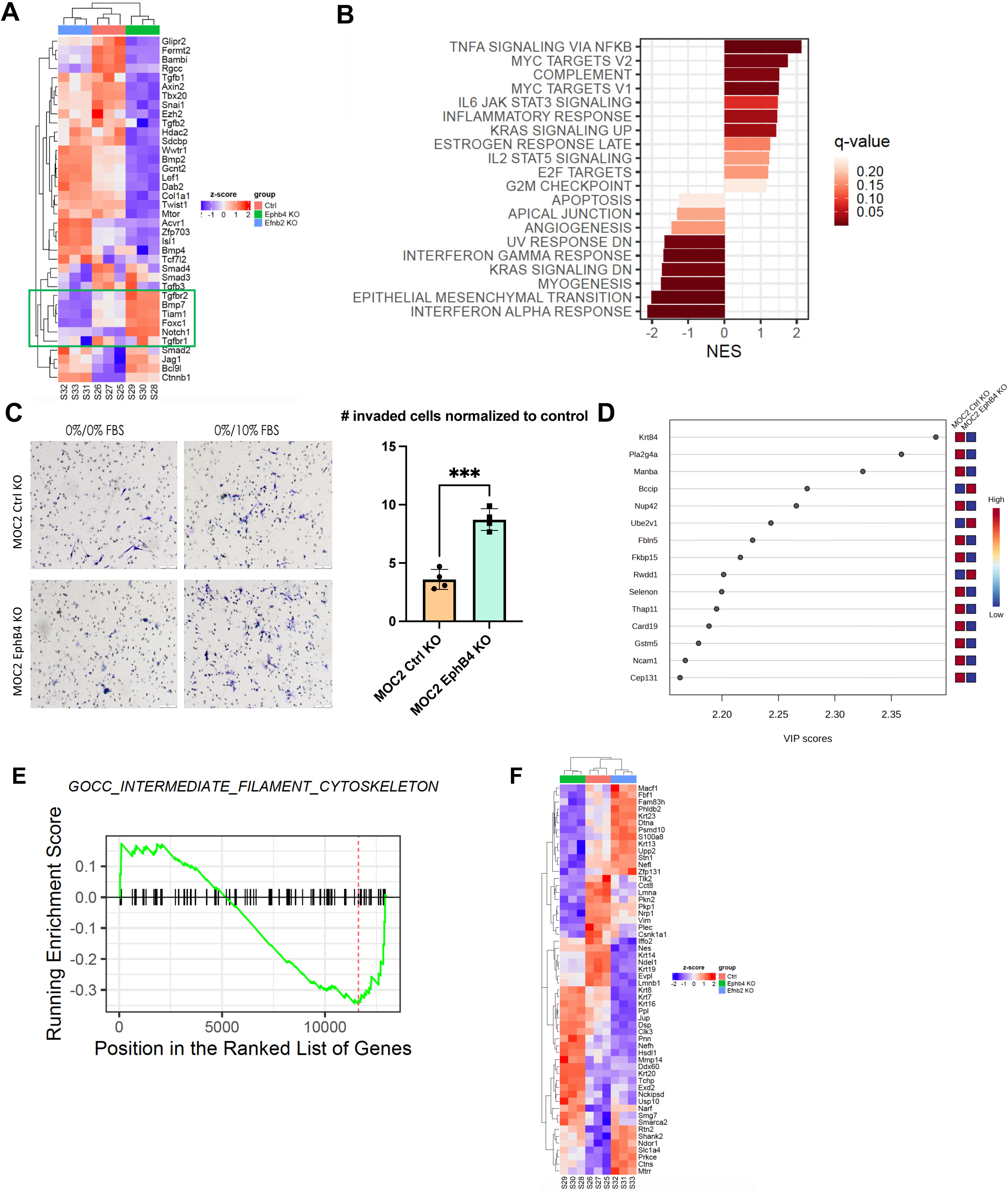
Loss of EphB4 in cancer cells induces protein dysregulation concomitant with increased metastatic capacity. (A) Expression of genes quantified using RNA-sequencing of MOC2 control, EphB4, and ephrinB2 (EFNB2) CRISPR knockout (KO) cell lines. (B) Hallmark pathways generated from RNA-sequencing of MOC2 control versus EphB4 shRNA cell lines. (C) Representative images and quantification for Boyden chamber invasion assay conducted on MOC2 control or EphB4 KO cell lines. (D) Variable importance in projection (VIP) score plots of mass spectrometry proteomics data conducted on MOC2 EphB4 KO vs Ctrl tumors displaying upregulation of BCCIP and Ube2v1 proteins in EphB4 KO tumors compared to control tumors. (E) GO and Reactome pathway enrichment analysis of EphB4 KO vs Ctrl MOC2 cancer cells. (F) Expression of genes associated with intermediate filament cytoskeleton quantified using RNA-sequencing of MOC2 control, EphB4, and ephrinB2 CRISPR knockout cell lines. For Boyden chamber quantification, comparison of invaded cells between the control and experimental group was done using a two-sided student’s *t*-test. Significance was determined if the *p*- value was <0.05*, <0.01**, <0.001***, and <0.0001****. *p*-values are indicated for the figures C \*\*\**p* = 0.0002.

Mass spectrometry proteomics analysis also revealed signatures consistent with enhanced metastatic potential in EphB4 knockdown cells (Figure 2D). The variable importance in projection (VIP) score plots derived from mass spectrometry-based proteomics data comparing MOC2 EphB4 knockout tumors to control tumors unveiled a significant upregulation of BCCIP and Ube2v1 proteins (Figure 2D). This finding holds substantial clinical relevance because overexpression of BCCIP has been associated with a poor prognosis and implicated in the facilitation of proliferation and migration in lung adenocarcinoma (Shi et al., 2021). In addition, Ube2v1, a member of the ubiquitin conjugating E2 enzyme variant proteins, plays a pivotal role in promoting EMT and metastasis by epigenetically suppressing autophagy at the transcriptional level (Shen et al., 2018). The observed upregulation of BCCIP and Ube2v1 in EphB4 knockout tumors suggests their potential involvement in tumorigenesis and metastatic progression. These data provide an association between the absence of EphB4 and cancer cell polarization towards a pro-metastatic phenotype.

To investigate the functional impact of EphB4 knockout, Gene ontology (GO) and Reactome pathway enrichment analyses were conducted for MOC2 cancer cells. Comparison of EphB4 knockout versus control cells revealed significant alterations in genes associated with the cytoskeleton (Figure 2E, F). Notably, EphB4 knockout cancer cells had low expression of Nefl, which has been shown to be associated with nodal spread and poor prognosis in breast cancer patients (Figure 2F) (Li et al., 2012). Nefl also induces apoptosis, suppresses growth, and decreases invasion of HNSCC cancer cells, offering potential mechanisms by which Nefl downregulation in EphB4 knockout cancer cells enhances their metastatic capacity (Huang et al., 2014). Additionally, EphB4 knockout cells had increased expression of matrix metallopeptidase 14, pinin, and keratins 7, 8, 16, and 20 (Figure 2F), genes associated with increased proliferation, EMT, and metastasis across multiple cancer types (Elazezy et al., 2021; Fang et al., 2017; Hosseinalizadeh et al., 2024; Niland et al., 2021; Tan et al., 2017; Wei et al., 2016). Collectively, these results suggest that EphB4 knockout induces molecular changes in MOC2 cancer cells, affecting regulation of the cytoskeleton, migration, and EMT. These changes may contribute to tumor progression and metastasis, highlighting the role of cancer cell EphB4 in regulating metastasis.

### Knockdown of EphB4 in cancer cells combined with radiation treatment promotes suppressive intratumoral immune populations

Building upon our RNAseq and mass spectrometry proteomics data (Figure 2), we used flow cytometry analysis to further understand the implications of cancer cell EphB4 knockdown *in vivo* (Figure 3A). Although the overall frequency of total CD4+ T-cells was the same in both EphB4 knockdown and control groups, we observed that EphB4 knockdown led to an increase in intratumoral Tregs (Figure 3B-D). We also observed upregulated markers associated with myeloid-derived suppressor cells (MDSCs) such as Ly6C+ Ly6G+ (Figure 3E), CD11b+Ly6G^high^Ly6G^low^ granulocytic MDSCs (Figure 3F), and CD11b+Ly6G^low^Ly6C^high^ monocytic MDSCs (Figure 3G). To determine if EphB4 knockdown in the cancer cell directly affects the CD4+ T cell state, we cocultured control or EphB4 shRNA cancer cells with CD4+ T cells and conducted flow cytometry (Figure 3H, Suppl. Figure 2). Our findings revealed that EphB4 knockdown in cancer cells polarizes CD4+ T cells towards the Treg phenotype (Figure 3J) and enhances Treg immunosuppression (Figure 3K).

**Figure 3:**
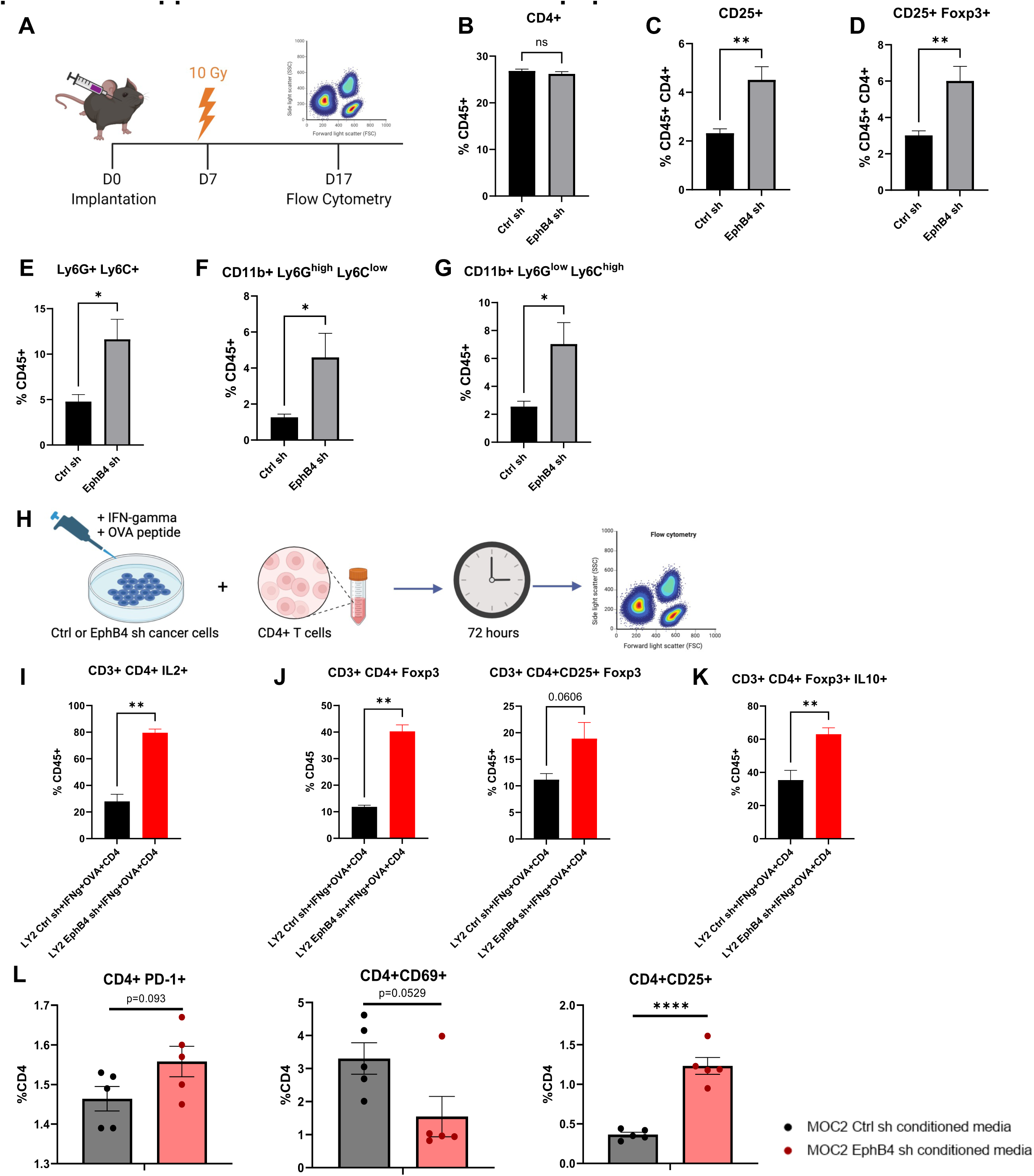
Knockdown of EphB4 in cancer cells combined with radiation treatment promotes suppressive intratumoral immune populations. (A) Experimental design for C57BL/6J mice implanted with MOC2 cancer cells. The buccal tumor received single dose of radiation treatment (10Gy) on day 7 after implantation. (B-G) Flow cytometric quantification of CD4+ T cells (B), regulatory T cells (Tregs) defined by CD4+ CD25+ (C) and CD4+ CD25+ Foxp3+ expression (D), myeloid-derived suppressor cells (MDSCs) defined by CD45+ Ly6G+ Ly6C+ (E), CD45+ Ly6G+ CD11b+ (F), and CD45+ Ly6C+ CD11b+ (G) in the tumor microenvironment (TME). H) Experimental design for coculture of LY2 Ctrl or EphB4 shRNA cancer cells with CD4+ T cells. Cancer cells were incubated with interferon-gamma (IFNg) for 48 hours, then OVA peptide overnight. CD4+ T cells from DO11.10 BALB/c mice were then cocultured with the cancer cells for 72 hours and subsequently harvested for flow cytometry. (I) Flow cytometry analysis displaying the quantification of IL2+ expressing CD4+ T cells. (J) Flow cytometry quantification of Tregs defined by CD4+ Foxp3+ and CD4+ CD25+ Foxp3+ expression. (K) Flow cytometry quantification of Treg immunosuppressive function defined by IL-10 expression. (L) Flow cytometry quantification of CD4+ T cells treated with conditioned media from MOC2 control or EphB4 shRNA cells showing PD-1, CD69, and CD25 expression. Comparison between control and experimental groups was done using a two-sided student’s *t*-test. Significance was determined if the *p*-value was <0.05*, <0.01**, <0.001***, and <0.0001****. *p*-values are indicated for the figures C ns *p* = 0.3636, D CD25+ \*\**p* = 0.0025; CD25+Foxp3+ \*\**p* = 0.0051, E \**p* = 0.0177, F \**p* = 0.0101, G \**p* = 0.0102, I \*\**p* = 0.0022, J \*\**p* = 0.0020, K \*\**p* = 0.0087, L \*\*\*\**p* < 0.0001.

Given the observed increase in Tregs with EphB4 knockdown in the cancer cell, we investigated whether CD4+ T cells within the TME differentiate into Tregs by treating CD4+ T cells with conditioned medium from MOC2 EphB4 knockdown cancer cells *in vitro* and conducting flow cytometry. We found a trend towards higher PD-1 expression in CD4+ T cells treated with EphB4 knockdown conditioned medium, indicative of greater exhaustion (Dong et al., 2019) (Figure 3L). EphB4 knockdown conditioned medium also yielded a trend towards decreased CD4+ T cell activation (defined by CD69 expression) and a significant increase in CD25+ Tregs (Figure 3L). These data are consistent with our previous work showing that EphB4 knockdown in cancer cells increases intratumoral IL-10 and G-CSF, thereby promoting Treg differentiation and immunosuppression (Bhatia et al., 2022).

In order to elucidate the underlying mechanism by which EphB4 shRNA influences Treg function, Tregs isolated from the spleen and lymph nodes of tumor-bearing mice (MOC2 control shRNA or EphB4 shRNA) were sorted and subjected to mass spectrometry proteomics analysis (Suppl. Figure 3A). Gene ontology (GO) processes revealed a concurrent upregulation of actin polymerization and depolymerization processes, leukocyte migration, and inflammatory response (Suppl. Figure 3B). Intriguingly, EphB4 shRNA-mediated knockdown also led to downregulation of pathways related to T helper 1 (Th1) and T helper 2 (Th2) cell differentiation as well as the mTOR signaling pathway, as indicated by KEGG pathway analysis (Suppl. Figure 3C). Remarkable decreases were also observed in microtubule organizing center function and TOR signaling by GO pathway analysis (Suppl. Figure 3D). Notably, the observed downregulation in mTOR signaling may contribute to Treg stabilization and maintenance of an immunosuppressive TME through the PI3K-Akt-mTOR signaling (Munn et al., 2018). These findings shed light on the potential molecular pathways through which EphB4 downregulation modulates Treg function, providing valuable insights into therapeutic targets that could be used for regulating immune responses in the context of HNSCC.

### Knockdown of ephrinB2 in the vasculature significantly reduces distant metastases

Since EphB4’s ligand ephrinB2 is expressed in blood vessels (Diehl et al., 2005; Wang et al., 2010), we sought to determine how vascular ephrinB2 affects metastatic spread of HNSCCs. Using the ephrinB2^fl/flTie2Cre^ (ephrinB2 KO) mouse model (Figure 4A) (Bhatia et al., 2022) implanted with MOC2 cancer cells, we found that deletion of vascular ephrinB2 coupled with RT significantly reduced local tumor growth compared to wild-type (WT) hosts (Figure 4B, Suppl. Figure 4). EphrinB2 knockout in vascular endothelial cells also significantly reduced the incidence of lung metastasis, leading to increased lung metastasis-free survival (Figure 4C, D). Additionally, our cell line RNA sequencing data showed lower expression of TGF beta receptors 1 and 2, NOTCH1, and FOXC1 in MOC2 ephrinB2 knockout cells compared to controls (Figure 2A). These findings collectively underscore the multicompartmental contributions of ephrinB2 in modulating HNSCC metastasis, highlighting it as a potential therapeutic target.

**Figure 4:**
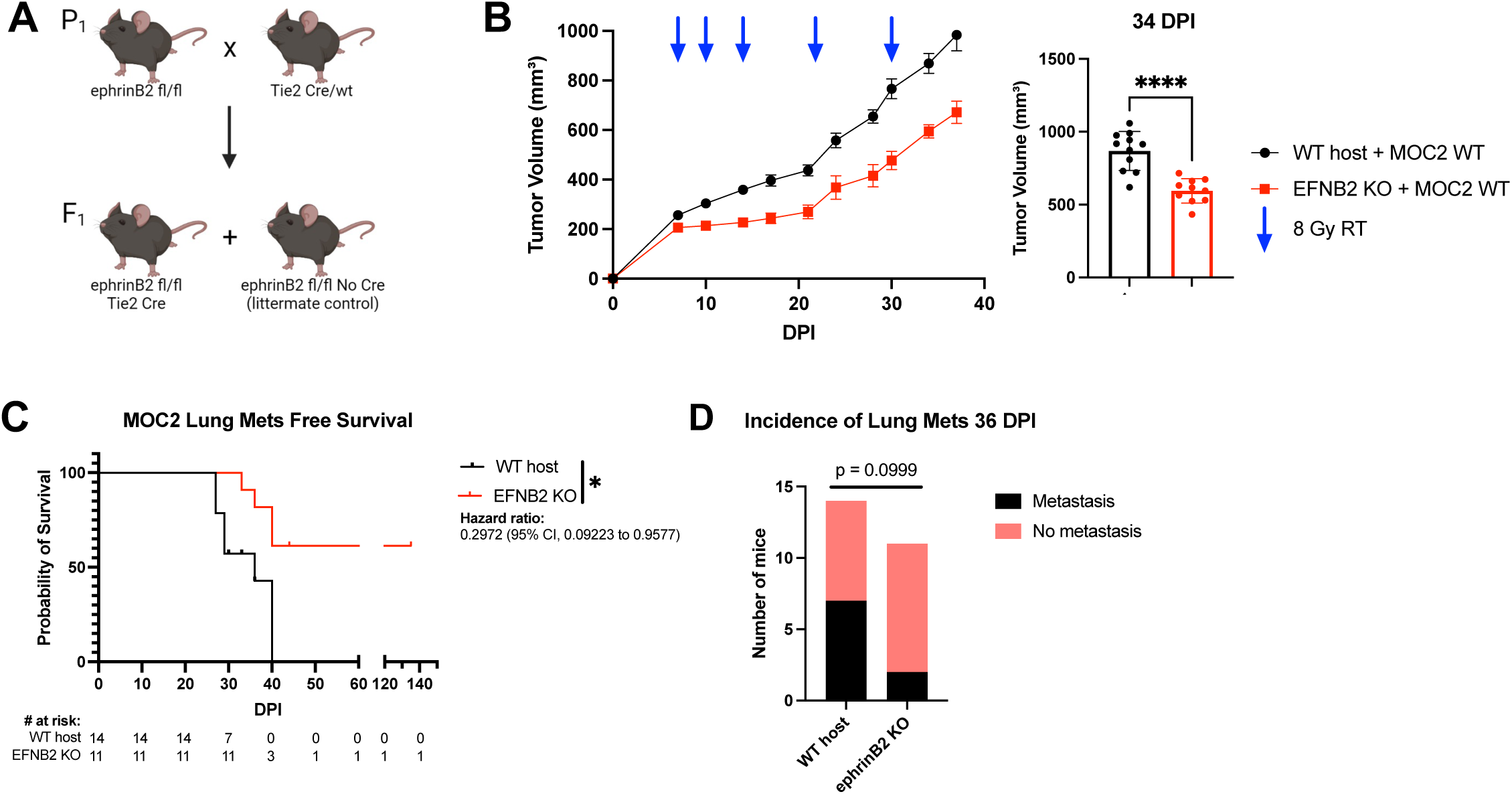
Knockout of ephrinB2 in the vasculature significantly reduces distant metastases. (A) Breeding strategy for creating mice with ephrinB2 (EFNB2) KO on vascular endothelial cells. (B) Average tumor volume curve and dot plot showing significantly smaller average tumor volume at 34 DPI in ephrinB2^fl/fl^Tie2^Cre^ mice implanted with MOC2 WT tumors compared to WT hosts. (C) Kaplan-Meier curves showing lung metastasis free survival of WT hosts and ephrinB2 KO mice. Numbers at risk indicate mice that were alive without metastases at specified timepoints. (D) Contingency table showing incidence of lung metastases detected by CT scans in ephrinB2 KO mice versus WT hosts by 36 DPI. Comparison of tumor volume between the control and experimental group was done using a two-sided student’s *t*-test. For Kaplan-Meier survival curves, significance was determined by a log-rank Mantel-Cox test. For contingency tables indicating the incidence of metastases, significance was determined by a Chi- square test. Significance was determined if the *p*-value was <0.05*, <0.01**, <0.001***, and <0.0001****. *p*-values are indicated for the figures B \*\*\*\**p* < 0.0001, C \**p* = 0.0172.

### Knockout of ephrinB2 in vascular endothelial cells enhances anti-tumor immune cell populations in the TME

To investigate how vascular ephrinB2 shapes the TME in HNSCC, we analyzed a publicly available scRNA-seq dataset of HNSCC patients (Puram et al., 2017b). We first confirmed expression of ephrinB2, SELP, and SELE in endothelial cells (Figure 5A). Next, gene expression profiles of endothelial cells derived from patients with high ephrinB2 expression were compared to those without ephrinB2 expression. Notably, these analyses show that endothelial cells with high ephrinB2 expression have lower levels of SELP and SELE, both of which are genes associated with promoting transendothelial migration of effector T cells (Figure 5B) (Gong et al., 2017).

**Figure 5:**
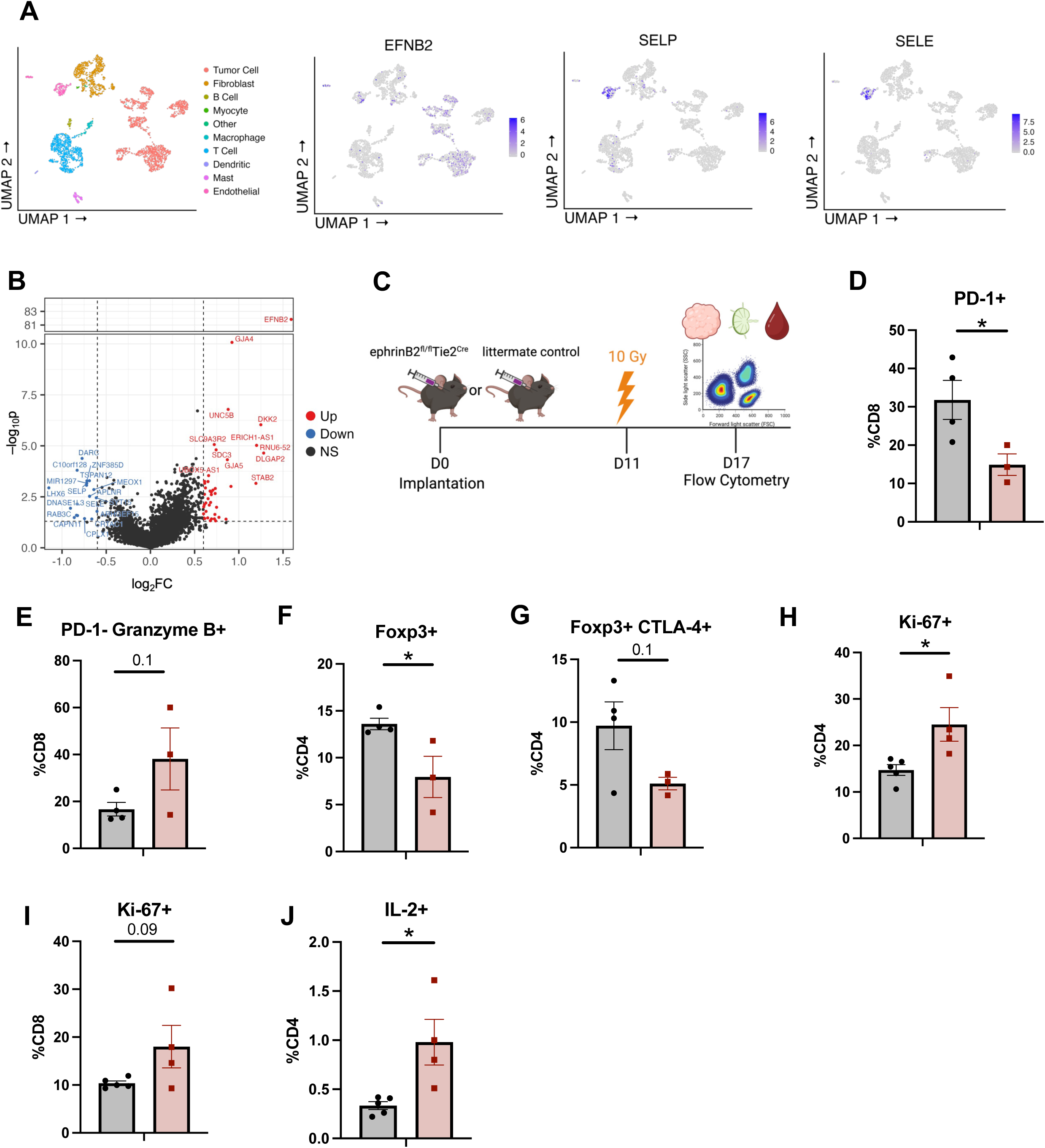
Knockout of ephrinB2 in vascular endothelial cells enhances anti-tumor immune cell populations in the TME. (A) UMAP plots showing expression of ephrinB2, SELP, and SELE in endothelial cells from human HNSCC single-cell sequencing. (B) Volcano plot showing differential gene expression of endothelial cells that do (red) or do not (blue) express ephrinB2 from human HNSCC single-cell sequencing. (C) Experimental design for flow cytometric analysis of MOC2 WT tumors implanted in ephrinB2 KO mice and littermate controls. (D-J) Flow cytometry analysis displaying quantification of PD-1 expressing CD8+ T cells (D), PD-1^-^ Granzyme B^+^ expressing CD8+ T cells (E), Tregs in the TME defined by Foxp3 expression (F), immunosuppressive Tregs defined by the addition of CTLA-4 expression (G), proliferating CD4+ T cells defined by Ki-67 expression in the draining lymph nodes (DLNs) (H), proliferating CD8+ T cells defined by Ki-67 expression in the DLNs (I), and IL-2 expressing CD4+ T cells in the DLNs (J). Comparison of differences between the control and experimental group was done using a two-sided student’s *t*-test. Significance was determined if the *p*- value was <0.05*, <0.01**, <0.001***, and <0.0001****. *p*-values are indicated for the figures D \**p* = 0.0466, F \**p* = 0.0358, H \**p* = 0.0247, J \**p* = 0.0177.

To further validate that vascular ephrinB2 modulates the immune cell composition of the TME after RT, we analyzed RNA sequencing of vascular endothelial cells isolated from C57BL/6 mice subjected to total body irradiation (5Gy x 1) (GSE141790). Six hours after RT, vascular endothelial cells were found to upregulate ephrinB2 RNA expression (Suppl. Figure 5A). EphrinB2 upregulation after RT was also confirmed in our MOC2 HNSCC tumor model, and ephrinB2-expressing vascular cells were observed to be more proliferative as determined by Ki- 67 expression (Suppl. Figure 5B, C). These data suggest that RT upregulates ephrinB2 in the vasculature, which may allow selective infiltration of immune cells.

We then conducted flow cytometry to examine how vascular ephrinB2 affects intratumoral immune cell composition in the context of RT (Figure 5C, Suppl. Figure 6). Within the TME, we found decreased PD-1 expression in CD8+ T cells, indicative of decreased exhaustion (Liu et al., 2021), along with increased granzyme B expression (Figure 5D, E). There was a significant decrease in Tregs and a trend towards decreased Treg expression of CTLA-4, suggesting decreased Treg-mediated immunosuppression (Sobhani et al., 2021) (Figure 5F, G).

Given the observed differences in the primary tumors, we sought to determine whether the changes seen in immune cells were limited to the TME due to tumor-targeted RT or whether global knockout of vascular ephrinB2 induced systemic changes. Analysis of the immune response in the blood and draining lymph nodes (DLNs) was conducted to answer this question (Suppl. Figure 7A, C). A trend towards higher CD8+ T cell granzyme B production was also observed in the circulation (Suppl. Figure 7B). Within the DLNs, there was increased proliferation of CD4+ and CD8+ T cells (Figure 5H, I), although no differences were found in their percentages (Suppl. Figure 7D, E). We also observed significantly higher CD4+ T cell activation and a Th1 phenotype, identified by increased IL-2 production (Figure 5J). Additionally, DLNs of ephrinB2 KO mice showed a trend towards increased dendritic cells (DCs), suggesting an environment where T cell priming may be more likely to occur (Suppl. Figure 7F). Similar to the TME, we observed a trend towards fewer Tregs in the DLNs (Suppl. Figure 7G). Collectively, these data suggest that vascular ephrinB2 may facilitate distant metastasis at least in part by dampening the systemic immune response.

To evaluate the effects of targeting ephrinB2 on lymphocyte extravasation into the TME, we conducted specialized flow cytometry experiments involving adoptive transfers of CD4+ T cells into tumor-bearing mice (Suppl. Figure 8A) (Estin et al., 2017; Thompson et al., 2020). Flow cytometry was conducted to quantify extravasation by distinguishing between intravascular and intratumoral lymphocytes (Suppl. Figure 8A) (Anderson et al., 2014; Estin et al., 2017; Thompson et al., 2020). This enabled us to extrapolate transendothelial migration from ratios of CD4+ T cells in the tumor versus in the vessels. Optimization studies were performed to determine that 1 hour following adoptive transfer was the appropriate time to allow CD4+ T cell extravasation within the MOC2 TME, as this timepoint preceded a rise in intratumoral CD4+ T cells (Suppl. Figure 8A, B). We found higher ratios of vessel to tumor CD4+ cells in ephrinB2 knockout mice, suggesting increased CD4+ T cell accumulation in the TME (Suppl. Figure 8C). Collectively, our data support inhibition of vascular ephrinB2 to enhance infiltration of anti-tumor immune populations and combat cancer progression.

### EFNB2-Fc-His and Fc-TNYL-RAW-GS reduce local and distant metastasis in C57BL/6 mice implanted with MOC2 cells

Since our current and previous data (Bhatia et al., 2022; Bhatia et al., 2019) suggest that it would be therapeutically desirable to maintain or activate EphB4 signaling in tumor cells while also inhibiting ephrinB2 reverse signaling in the vasculature, we examined the *in vivo* effects of a dimeric ephrinB2-Fc fusion protein in our MOC2 mouse model of metastasis. Ephrin Fc fusion proteins bind to the ephrin-binding pocket in the ligand-binding domain of Eph receptors to activate receptor forward signaling, inhibit Eph receptor-ephrin interaction, and thus inhibit ephrin reverse signaling (Barquilla et al., 2016). We also examined the effects of Fc-TNYL-RAW-GS, a version of the EphB4-targeting TNYL-RAW peptide dimerized by fusion of its N-terminus to Fc, since N-terminal dimerization has been shown to transform TNYL-RAW from an antagonist to an agonist (Fan et al., 2022; Mitchell Koolpe et al., 2005). The TNYL-RAW peptide, like ephrinB2, targets the ephrin-binding pocket of EphB4 (Mitchell Koolpe et al., 2005) and inhibits ephrinB2 reverse signaling (Noberini et al., 2011). However, TNYL-RAW is more specific since it only binds to EphB4 among the Eph receptors (Mitchell Koolpe et al., 2005), whereas ephrinB2 also binds to the other EphB receptors and to EphA4 (Elena B. Pasquale, 2004).

We used hydrodynamic injection of plasmids encoding EFNB2-Fc-His or Fc-TNYL-RAW-GS to transfect mouse liver cells and the Sleeping Beauty (SB) system to achieve prolonged protein expression and secretion of the Fc proteins by liver cells into the mouse circulation (Figure 6A) (Aronovich et al., 2007; Liang et al., 2018). Administration of EFNB2-His-Fc or Fc-TNYL-RAW-GS in conjunction with RT reduced local tumor growth and improved overall mouse survival (Figure 6B-D, Suppl. Figure 9). Additionally, EFNB2-Fc-His and Fc-TNYL-RAW-GS increased lung metastasis-free survival compared to controls (Figure 6E). Notably, by the time half of the control mice had lung metastases detected by CT scans, mice treated with EFNB2-Fc-His had no incidence of metastases (Figure 6F).

**Figure 6:**
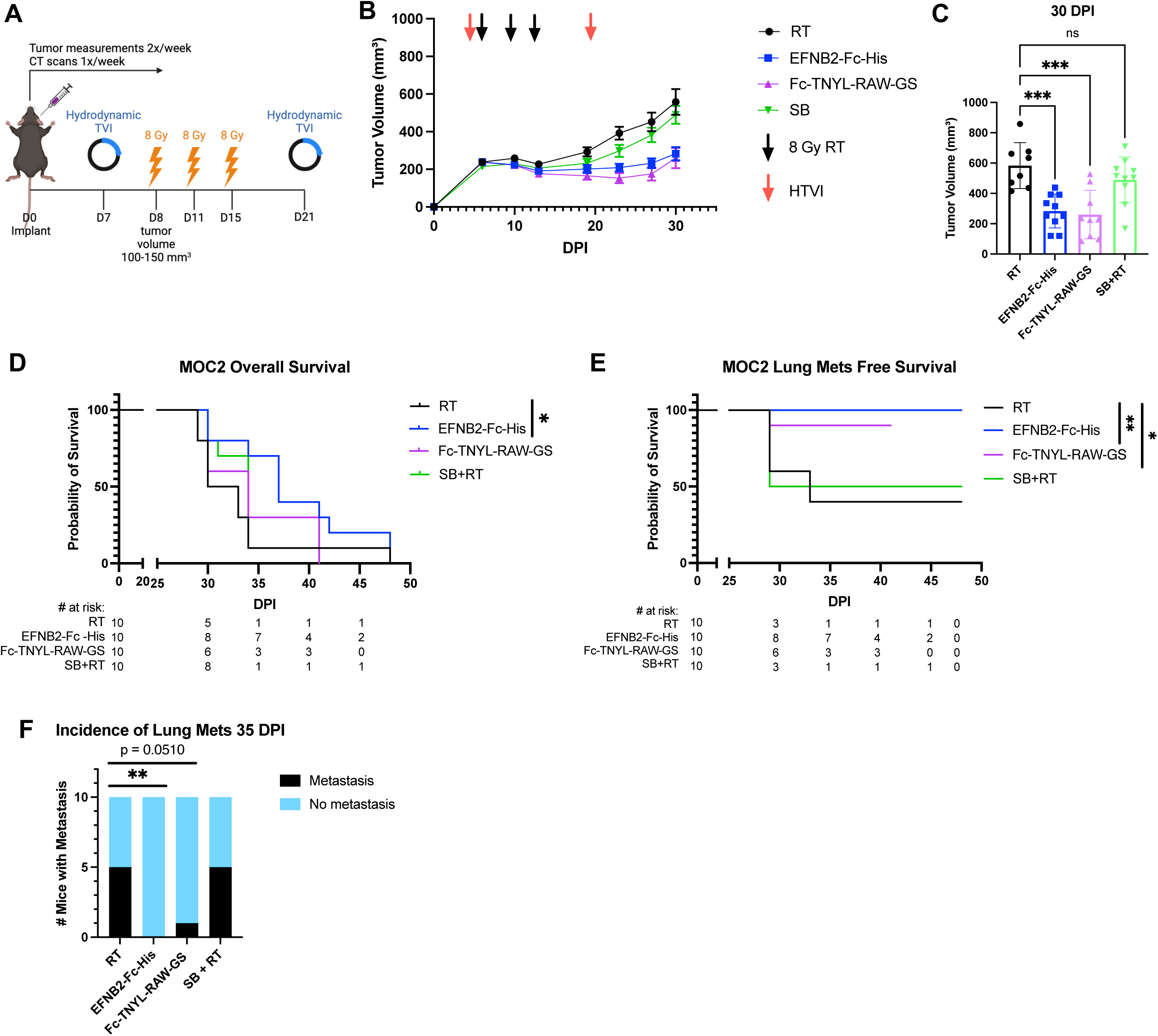
EFNB2-Fc-His and Fc-TNYL-RAW-GS reduce local tumor growth and distant metastasis in C57BL/6 mice implanted with MOC2 cells. (A) Experimental design for C57BL/6J mice implanted with MOC2 WT cancer cells. Hydrodynamic tail vein injections (HTVI) and radiation therapy (8 Gy) were administered as indicated. Mice treated with RT alone or Sleeping Beauty (SB) + RT served as controls. (B) Average tumor volume curve comparing effects of different plasmids on local tumor growth. (C) Dot plot showing the effects of different plasmids on tumor volume 30 DPI. (D-E) Kaplan-Meier curves showing therapeutic effects of different plasmids on overall survival (D) and lung metastasis free survival (E) in MOC2 WT implanted C57BL/6 mice. For Kaplan-Meier curves showing lung metastasis free survival, numbers at risk indicate mice that were alive without metastases at specified timepoints. (F) Contingency table quantifying the incidence of lung metastasis detected by CT scans in MOC2 WT implanted mice by 35 DPI. Comparison of tumor volume between the control and experimental groups was done using a Dunnett post hoc test after one-way ANOVA was performed. For Kaplan-Meier survival curves, significance was determined by a log-rank Mantel-Cox test. For contingency tables indicating the incidence of metastases, significance was determined by a Chi-square test. Significance was determined if the *p*-value was <0.05*, <0.01**, <0.001***, and <0.0001****. *p*-values are indicated for the figures C RT vs EFNB2-Fc-His \*\*\**p* = 0.0003; RT vs Fc-TNYL-RAW-GS \*\*\**p* = 0.0002; RT vs SB+RT ns *p* = 0.3757, D \**p* = 0.0485, E RT vs EFNB2-Fc-His \*\**p* = 0.0075; RT vs Fc-TNYL-RAW-GS \**p* = 0.0488, F \*\**p* = 0.0098.

### EFNB2-Fc-His activates EphB4 without concurrent activation EFNB2 while Fc-TNYL-RAW-GS activates both EphB4 and EFNB2

To validate the mechanisms through which EFNB2-Fc-His and Fc-TNYL-RAW-GS reduce local tumor growth and the formation of distant metastases, we repeated hydrodynamic injections of these plasmids and included EFNB2-Fc for comparison (Figure 7A). Concordant with our previous data, EFNB2-Fc, EFNB2-Fc-His, and Fc-TNYL-RAW-GS all significantly reduced local tumor growth while EFNB2-Fc-His was the only treatment that yielded a significant improvement in overall survival (Figure 7B-D). Tumors were harvested and processed for western blotting, then probed for expression of EphB4, phospho-EphB4 (pEphB4), EFNB2, and phospho-EFNB2 (pEFNB2) (Figure 7E). Comparing the ratios of relative protein expression of pEphB4/EphB4 between control and experimental tumors, EFNB2-Fc and EFNB2-Fc-His significantly activated EphB4 while Fc-TNYL-RAW-GS trended towards EphB4 activation (Figure 7F). Additionally, the ratios of relative protein expression of pEFNB2/EFNB2 in control and experimental tumors demonstrated that EFNB2-Fc-His significantly reduces activation of EFNB2 while Fc-TNYL-RAW-GS significantly activates EFNB2 (Figure 7F).

**Figure 7:**
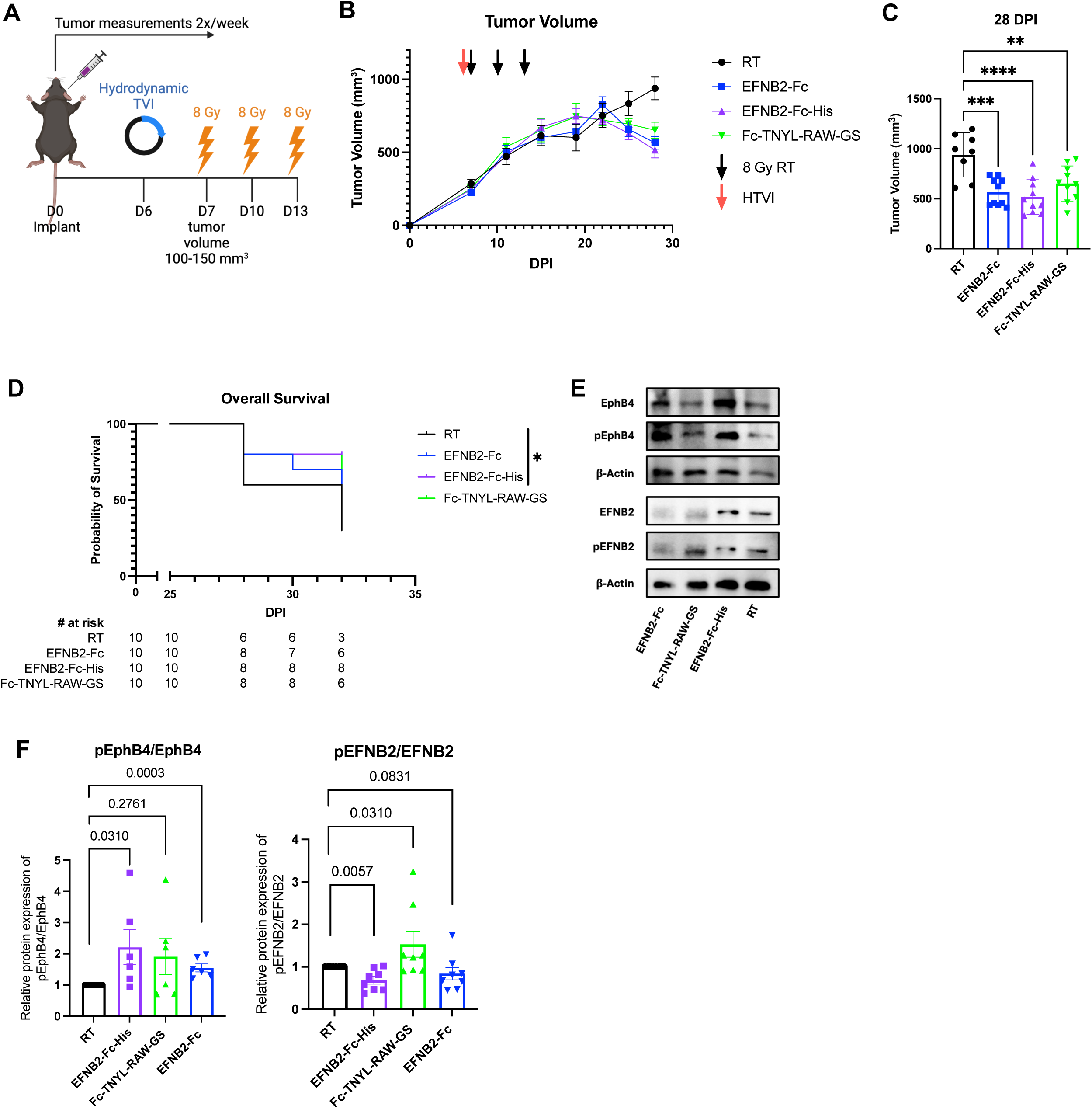
EphrinB2-Fc-His activates EphB4 without concurrent activation of EFNB2 while Fc- TNYL-RAW-GS activates both EphB4 and EFNB2. (A) Experimental design for C57BL/6J mice implanted with MOC2 WT cancer cells. Hydrodynamic tail vein injections (HTVI) and radiation therapy (8 Gy) were administered as indicated. Mice treated with RT alone served as controls. (B) Average tumor volume curve comparing effects of different plasmids on local tumor growth. (C) Dot plot showing the effects of different plasmids on tumor volume 28 DPI. (D) Kaplan-Meier curves showing therapeutic effects of different plasmids on overall survival in MOC2 WT implanted C57BL/6 mice. (E) Western blots showing protein expression of EphB4, phospho-EphB4 (pEphB4), EFNB2, phospho-EFNB2 (pEFNB2), and beta-actin in control and experimental tumors. (F) Dot plots showing ratios of protein expression of pEphB4/EphB4 and pEFNB2/EFNB2 in control and experimental tumors. Protein bands were quantified using Image lab and ImageJ software. Comparison of tumor volume and relative protein expression between the control and experimental groups was done using Mann-Whitney tests. Significance was determined if the *p*-value was <0.05*, <0.01**, <0.001***, and <0.0001****. *p*-values are indicated for the figures C RT vs EphrinB2-Fc \*\*\**p* = 0.0002; RT vs EFNB2-Fc-His \*\*\*\**p* < 0.0001; RT vs Fc-TNYL-RAW-GS \*\**p* = 0.0044, D \**p* = 0.0389.

To verify successful *in vivo* expression of Fc fusion proteins, serum was collected 26 days following hydrodynamic injection and probed by immunoblotting with anti-human Fc antibodies. Fc fusion proteins were detected in the serum of mice injected with EFNB2-Fc-His and Fc-TNYL-RAW-GS (Suppl. Figure 10A). Additionally, an ELISA was performed to measure EphB4 phosphorylation in PC3 prostate cancer cells and EphB4 stably expressing HEK293 cells following treatment with commercially available ephrinB2-Fc (ephrinB2-Fc R&D), EFNB2-Fc-His, and Fc-TNYL-RAW-GS (Suppl. Figure 10B). Compared to the Fc control, all three treatments resulted in phosphorylation of EphB4 (Suppl. Figure 10B).

Our data suggest that EphB4 acts as a tumor suppressor by inhibiting both the growth of primary tumors and the formation of metastases. In contrast, vascular EFNB2 acts as a promoter of local tumor growth and distant metastases. Our phopho-western blot data support that activating EphB4 while simultaneously avoiding activation of EFNB2 using various therapeutic strategies such as the EFNB2-Fc-His plasmid provides maximal benefit to reducing local tumor growth and distant metastases. Taken together, our results emphasizes the role of EphB4 as a suppressor of cancer cells’ intrinsic metastatic capacity, which can be amplified by agents that activate EphB4. Our data also underscore the therapeutic benefit of inhibiting ephrinB2 signaling in vascular endothelial cells and systemically. For the first time, we also provide therapetuic approaches that can activate EphB4 and inhibit ephrinB2 therefore serving as key translational strategies.

## Discussion

The EphB4-ephrinB2 signaling axis is aberrantly expressed in a variety of cancers, highlighting it as a promising therapeutic target (Alam et al., 2008; Broggini et al., 2020; Piffko et al., 2022; Sharma et al., 2015; Xu et al., 2023). While some studies implicate EphB4-ephrinB2 signaling as a pro-metastatic driver (Héroult et al., 2010; Sharma et al., 2015), others show the opposite (Broggini et al., 2020; Piffko et al., 2021). Contributing to the complexity is the fact that EphB4 and ephrinB2 can have independent, dichotomous effects on primary tumor growth and metastasis (Bhatia et al., 2022; Li et al., 2019; Noren & Pasquale, 2007; Sasabe et al., 2017). Consequently, it is increasingly apparent that context is key to deciphering the role of this receptor-ligand pair, both in terms of cancer type and cellular compartment (Pasquale, 2010; Pergaris et al., 2021). However, many studies on EphB4-ephrinB2 signaling in HNSCC metastasis have been limited to descriptive correlations and/or heavy reliance on *in vitro* experiments (Masood et al., 2006; Sasabe et al., 2017; Sinha et al., 2006; Yi et al., 2021). Our study provides novel insight into the intricate dynamics of EphB4-ephrinB2 signaling in HNSCC metastasis *in vitro* and *in vivo*.

Specifically, we illuminate the role of EphB4 knockdown in cancer cells, shedding light on its impact on metastatic potential and the immune microenvironment. Our findings indicate that knockdown of EphB4 in cancer cells leads to increased metastasis in two models of HPV-unrelated HNSCC due to multiple factors. Our mass spectrometry proteomics analysis of EphB4 knockdown in MOC2 cells revealed dysregulation of proteins involved in cell division, which may enhance the intrinsic capacity of cancer cells to metastasize. Our RNA sequencing data also show significant alterations in immune and tumor-related pathways, including downregulation of Th1/Th2 cell differentiation and mTOR signaling. These findings suggest that EphB4 expressed in cancer cells modulates Treg function. Notably, we observed an augmentation in the prevalence of immunosuppressive Tregs and myeloid cells *in vivo*, potentially contributing to tumor immune evasion and providing insights into cancer immunotherapy targets within the TME. Our data suggest that this increase in intratumoral Tregs is driven by two processes – direct conversion of CD4+ T cells and increased infiltration of Tregs into the tumor.

Our lab and others have identified the vasculature as a main compartment of ephrinB2 activity in multiple cancer types including HNSCC (Bhatia et al., 2022; Broggini et al., 2022; Héroult et al., 2010; Noren et al., 2004). We previously reported that ephrinB2 knockout in the vascular endothelium in the absence of RT decreases local tumor growth and the abundance of Tregs in the TME (Bhatia et al., 2022). In this study, we sought to examine vascular ephrinB2 from a metastatic angle and incorporated RT due to the ability of RT to enhance anti-tumor immunity (Bickett et al., 2021; Darragh et al., 2022; M. W. Knitz et al., 2021; Oweida et al., 2018; Oweida et al., 2017; Oweida, Darragh, et al., 2019) and its use as a standard-of-care therapy for HNSCC patients (Pfister et al., 2020; Xing et al., 2021). Our characterizations of ephrinB2-expressing endothelial cells and extravasation data suggest that vascular ephrinB2 may modulate trafficking of lymphocytes through regulation of adhesion molecules. We observed that RT coupled with deletion of vascular ephrinB2 decreases metastasis in a preclinical mouse model of HNSCC. Additionally, our data show that vascular ephrinB2 loss combined with RT enhanced systemic CD4+ and CD8+ T cell activation, trended towards decreased Treg immunosuppression, and reduced CD4+ T cell entry into the tumor. As a result, concurrent RT and inhibition of vascular ephrinB2 is a potential therapeutic avenue to mitigate both local and distant HNSCC progression.

The promiscuous and compensatory nature of Eph receptors and ephrins (E. B. Pasquale, 2004; Pasquale, 2010; Pasquale, 2024) poses a challenge in efforts to specifically target metastasis-promoting components while maintaining protective signaling components. Our findings show that EphB4 and ephrinB2 have opposing effects on metastasis in the HPV-unrelated HSNCC MOC2 and LY2 models. Clinical trials incorporating broad receptor pan-targeted tyrosine kinase inhibitors that may affect Eph-ephrin signaling, including erlotinib, dasatinib, and cetuximab, have largely failed to provide meaningful clinical benefit due to limited efficacy and resistance (Hermida-Prado et al., 2019; Kang et al., 2023; Li et al., 2023; Sola et al., 2019). Given the structural similarities and signaling cascades (PI3K, mTOR, MAPK, JAK/STAT, etc.) shared among receptor tyrosine kinases (Du & Lovly, 2018; Regad, 2015), inhibitors targeting them also pose the risk of off-target effects, including potential inhibition of the Eph receptor and ephrin gene families. Consequently, exploration of more targeted therapeutics is needed to improve patient response.

A notable finding in this study is the capacity of EphrinB2-Fc-His and Fc-TNYL-RAW-GS to decrease local tumor growth and the incidence of lung metastases in a preclinical model of HNSCC. Given their high specificity for their targets (EphB receptors and EphA4 for EFNB2- Fc-His or EphB4 for Fc-TNYL-RAW-GS), the two Fc fusion plasmids are promising agents to combat HNSCC metastasis and circumvent off-target effects of traditional receptor tyrosine kinase inhibitors (Duggineni et al., 2013; Fan et al., 2022; M. Koolpe et al., 2005; Noberini et al., 2011). The benefits observed in our preclinical models open the door to further exploration of other Fc fusion proteins targeting Eph receptors and ephrins depending on their expression levels and presumed effects in certain cancer types. In light of our findings, one potential therapy to abrogate HNSCC metastasis would be an EphB4 agonist that targets the ephrin-binding pocket to activate EphB4 and inhibit ephrinB2.

## Materials and Methods

### Cell Culture and Reagents

Murine HNSCC tumor cell lines MOC2 and LY2 were used for all *in vivo* studies. The MOC2 cell line is derived from a C57BL/6 mouse with squamous cell carcinoma of the oral cavity that was exposed to DMBA for 25 weeks. The LY2 cell line was obtained from Dr. Nadarajah Vigneswaran (University of Texas Health Science Center, Houston, Texas, USA). The LY2 cell line was isolated from lymph node metastases that developed in BALB/c mice after inoculation of PAM 212 squamous cell carcinoma cells (Chen et al., 1997). All cell lines were cultured at 37°C and 5% CO_2_. The LY2 cell lines were cultured in DMEM-F12 media supplemented with 10% fetal bovine serum, 2% Primocin, and 1% Fungin (InvivoGen, San Diego, California, USA). The MOC2 cell line was cultured in DMEM-F12:IMDM (2:1) supplemented with 2% Primocin and 1% Fungin (InvivoGen, San Diego, California, USA), 1.75ng EGF, 20ng hydrocortisone per 500mL of media, and 0.1% insulin (Sigma Aldrich, St. Louis, Missouri, USA). HEK293AD human embryonic cells (#240085, Agilent, Santa Clara, California, USA) and PC3 prostate cancer cells (#CRL-1435, ATCC) were cultured in Dulbecco’s Modified Eagle Medium (DMEM; #10-013-CV, Corning, Corning, New York, USA) and RPMI 1640 medium(#11875093, ThermoFisher Scientific, Waltham, Massachusetts, USA), supplemented with 10% fetal bovine serum and 1% antimycotics and antibiotics (#A5955, Sigma-Aldrich, St. Louis, Missouri, USA). HEK293AD cells were stably transfected with pLVX-IRES-Neo encoding full-length human EphB4 with an N-terminal FLAG tag using Lipofectamine 2000 reagent according to the manufacturer’s instruction (#11668019, ThermoFisher Scientific). Forty-eight hours after transfection, the cells were selected with 1 mg/ml G418 (#10131035, ThermoFisher Scientific) for 15 days to generate stably transfected cells. T cells were cultured in RPMI media supplemented with recombinant human IL-2 (Biological Resources Branch National Cancer Institute, Frederick, Maryland, USA), 10% fetal bovine serum, 50nM 2-mercaptoethanol, 1X MEM non-essential amino acids, 1X penicillin-streptomycin, 1M HEPES, and 100mM sodium pyruvate (Gibco, Billings, Montana, USA).

### Fc fusion proteins

Commercial purified mouse ephrinB2-Fc was purchased from R&D Systems (#496-EB-200, which contains a C-terminal His tag). The ephrinB2-Fc-His plasmid was obtained by cloning the cDNA encoding the mouse ephrinB2 extracellular region (residues 3-227), a GSGDP linker, the Fc portion of human IgG1, a glycine, and a His tag into the PT3-EF1a-C-Myc plasmid. The Fc- TNYL-RAW-GS plasmid was obtained by cloning a cDNA encoding the CD5 signal peptide followed by the Fc portion of human IgG1 and the sequence ARTNYLFSPNGPIARAWGS, with the underlined sequence representing the TNYL-RAW peptide {Koolpe, 2005 #98}. The Fc control plasmid was obtained by cloning a cDNA encoding the CD5 signal peptide followed by the Fc portion of human IgG1. The PT3-EF1a-C-Myc and pCMV/SB11 plasmids were kindly provided by Dr. Gen-Sheng Feng (University of California, San Diego) {Chen, 2021 #139}.

Purified ephrinB2-Fc-His and Fc-TNYL-RAW-GS proteins for cell stimulation were generated by transiently transfecting HEK293 cells with the respective plasmids and the pCMV/SB11 plasmid at a 1:10 ratio. Cells were passaged into a larger plate the next day. Upon reaching 70% confluence, the culture medium was replaced with fresh Opti-MEM (#31985088, ThermoFisher Scientific) after washing the cells twice with warm DPBS (#21-030-CV, Corning). After three days, the culture medium was collected, and HEPES, pH 7.5 was added to a final concentration of 10 mM before centrifugation at 1,000 g for 10 minutes to remove cell debris. The resulting supernatant was incubated with GammaBind Plus Sepharose beads (#17088602, Cytiva, Marlborough, Massachusetts, USA) overnight at 4C. Beads were washed with cold PBS once before bound Fc fusion proteins were eluted with 0.1 M glycine HCL, pH 2.5, and the low pH in the eluates was neutralized with 1 M Tris HCl buffer, pH 7.5.

### Animal Tumor Models

C57BL/6 mice were purchased from the Jax Labs, Bar Harbor, Maine, USA. In vivo orthotopic HNSCC tumor models were established as previously described (Darragh et al., 2022; Oweida, Bhatia, et al., 2019). Deletion of ephrinB2 from the vasculature of ephrinB2fl/flTie2Cre mice was done as previously described (Bhatia et al., 2022). For buccal tumor implantations, 1 x 10^5^ MOC2 cells per 50ul of serum-free cell media were prepared. For floor of mouth implantations, 1 x 10^5^ LY2 cells per 50uL of serum-free cell media were prepared. A 1:1 mixture of cells and Matrigel (10mg/mL, BD Biosciences, San Jose, California, USA) at a volume of 100ul was injected into the right buccal mucosa of the mice (MOC2) or floor of mouth (LY2). Radiation treatment was given as described below when tumors reached ∼100-200mm^3^ (MOC2) or ∼150- 300mm^3^ (LY2). Tumor size was measured twice weekly using digital calipers, and tumor volume was estimated using the equation V=AxB2/2, where A is the longest diameter of the tumor and B is the shortest. Computed tomography scans were performed once a week as described below. Mice were euthanized when the mice reached study end qualifications approved by the Institutional Animal Care and Use Committee (IACUC). Mice were euthanized under CO_2_, and intracardiac blood collection was performed. Blood was placed in serum collection tubes (BD, Franklin Lakes, New Jersey, USA) and centrifuged at 6000 rpm for 2 minutes to collect the serum supernatant. Tumors were harvested and flash frozen in liquid nitrogen, while lung tissues were harvested and placed in 10% formalin for further processing. Both sera and tumors samples were stored in -80°C for subsequent analysis.

For studies using ephrinB2-Fc and Fc-TNYL-RAW-GS plasmids, C57BL/6J mice were initially implanted with 100k MOC2 cancer cells. 7 days later, mice were randomized into different groups including a control group treated with radiation therapy alone, a control group transfected with 1µg of Sleeping Beauty transposase plasmid (SB), and experimental groups transfected with 1µg of SB in addition to 20 µg of EFNB2-Fc or Fc-TNYL-RAW-GS plasmid. Sleeping Beauty transposase plasmid was used to promote cDNA integration into mouse liver genomic DNA. Liver transfection was performed by hydrodynamic tail vein injection as described previously (S. Bhatia et al., 2019). A repeat liver transfection was performed 3 weeks post-tumor implantation. All animal protocols used in this study were approved by IACUC of the University of Colorado, Denver.

### Irradiation and Computed Tomography Scans

Mice were anesthetized using isoflurane before and during the procedure. The mice were then placed in an X-RAD image guided small animal irradiator (Precision X-Ray, Bradford, Connecticut, USA). For tumor irradiation, the irradiation field was determined using fluoroscopy for each mouse. After the field was established, the mice were irradiated with 225kVp/20mA with a copper filter, which corresponds to 5.6 Gy/min for 98 seconds (8Gy) or 123 seconds (10Gy). For computed tomography scans, mice were scanned using an aluminum filter. For MOC2 metastasis studies, 3D lung contouring was generated using ITK-Snap. For flow studies, mice received one fraction of 10 Gy. For MOC2 metastasis studies, three fractions of 8 Gy were administered to the buccal tumor. For LY2 floor of mouth studies, three fractions of 8 Gy were administered to the tumor. In some studies utilizing the MOC2 cell line, tumors were given an additional dose of 8 Gy RT as they approached 500mm^3^ in volume to reduce local tumor burden and prolong the study. For intravital studies, mice received one dose of 10Gy 12 to 14 days after implantation.

### Flow Cytometry

Tumors and draining lymph nodes (DLNs) were harvested from mice and immediately put on ice in Hank’s Balanced Salt Solution (HBSS). To facilitate single cell suspension of lymphocytes from tumors, the tumors were minced and then incubated for 30 minutes at 37 °C with 200U of Collagenase III (Worthington, Lakewood, New Jersey, USA). Both tumors and draining lymph nodes were then filtered through 70µm nylon filters. The tumors were then spun down at 400g for 5 minutes and resuspended in 3 mL of Red Blood Cell Lysis Buffer for 3 minutes. After 3 minutes, 6 mL of HBSS was added and all the samples were spun down again at 400g for 5 minutes. Blood was taken using an intracardiac puncture. The blood was spun down for 10 minutes at 400g before resuspending in 3mL of Red Blood Cell Lysis Buffer for 3 minutes. After 3 minutes, 6 mL of HNSCC was added to the sample and the samples were spun down at 400g for 5 minutes. The samples were aspirated and then were plated in 24 well plates in 1 mL of RPMI with 0.1% monensin to prevent cytokine release by the golgi apparatus and 0.2% brefeldin A with cell stim cocktail of PMA and ionomycin to simulate cytokine production and transport in the cells. The cells were incubated in the stim media for 4 hours at 37 C. The plates were then spun down and the samples were resuspended in 200µl of Fc Block (anti-CD16/CD32 antibody, Tonbo Biosciences, San Diego, California, USA) for 20 minutes at room temperature. The samples were then plated into a 96 well plate and spun down at 400g for 5 minutes. The samples were then resuspended in 100µl of PBS containing 5µl of live/dead aqua viability stain (Invitrogen, Carlsbad, California, USA). The samples were incubated away from light for 15 minutes at room temperature. The samples were then spun down and resuspended in a mixture of the extracellular antibodies in brilliant stain buffer (BD Biosciences, Franklin Lakes, New Jersey, USA). The cells were incubated for 30 minutes at room temperature. After incubated the cells were washed twice with FA3 buffer. The cells were then incubated overnight with the Foxp3 transcription factor staining kit (eBioscience, San Diego, California, USA). The next day the cells were spun down and washed twice as per the kit instructions. The cells were then stained with the intracellular antibodies in brilliant stain buffer for 30 minutes at room temperature. The cells were then washed twice with PBS. Cells were re-suspended and then fun on an Aurora Spectral Flow Cytometer (Cytek Biosciences, Fremont, California, USA) at the Barbara Davis Center Diabetes Research Center Cell and Tissue Analysis Core.

The following antibodies were used in these studies: PerCP-CD45 (Clone: 30-F11 Biolegend, San Diego, California, USA), PerCP-Cy5.5- CD3 (Clone: 17A2 Biolegend), BUV805-CD3 (Clone: 17A2 BD Biosciences), SB436-CD4 (Clone: GK1.5 Invitrogen), ef450-CD4 (Clone: GK1.5 Invitrogen), BUV496-CD4 (Clone GK1.5, BD 612952), BB515-CD8 (Clone: 53-6.7 BD Biosciences), BV570-CD44 (Clone: IM7, Biolegend 103037), AF532-Foxp3 (Clone: FJK-16s Invitrogen), ef660-CD11b (Clone: M1/70 Invitrogen), BUV661 (Clone: M1/70 BD Biosciences), BUV 711-CD25 (Clone: PC61, Biolegend 102049), NovaFluor yellow 590-CD11b (Clone: ICRF44, eBioscience), eFluor 450- LY6C (Clone; HK1.4, eBioscience), BV650-LY6C (Clone: HK1.4 Biolegend), BV570-LY6C (Clone: HK1.4 Biolegend), BV605-LY6C (Clone: HK1.4 Biolegend), PerCP Cy5.5-LY6G (Clone 1A8, Biolegend 127615), BV421-LY6G (Clone: 1A8 Biolegend), BV650-LY6G (Clone: 1A8 BD Biosciences), BV605-LY6G (Clone: 1A8 Biolegend), AF594-ephrinB2 (1° Catalogue number AF496 R&D; 2° Ref A21468), BV480- F4/80 (Clone: T45-2342 BD Biosciences), AF-647-IL2 (Clone: JES6-5H4, Biolegend 503814), BV605-IL-2 (Clone: JES6-5H4 Biolegend), PE-IL-2 (Clone: JES6-5H4 Biolegend), BV 421-IL-10 (Clone: JES5-16E3, Biolegend 505021), BUV395-PD-1 (Clone: J43 BD Biosciences), APC- R700-CTLA-4 (Clone: UC10-4F10-11 BD Biosciences), APC-Fire 750-Granzyme B (Clone: QA16A02 Biolegend), APC-iNOS (Clone: CXNFT Invitrogen), PerCP-eFlour 710-Arginase 1 (Clone: A1exF5 Invitrogen), APC-eFluor 780-Ki-67 (Clone: SoIA15, Invitrogen 47-5698-82), BV480-Ki-67 (Clone: B56 BD Biosciences), BV786-Ki-67 (Clone: B56 BD Biosciences), BV805-MHCII (Clone: M5/114.15.2 BD Biosciences), PE-Dazzle 594-MHCII (Clone: M5/114.15.2 Biolegend), PE-Cy7-CD31 (Clone: 390 Biolegend), SB436-PDPN (Clone: eBio.1.198.1.10 Invitrogen), Live Dead Aqua (Invitrogen).

The following antibodies were used in intravital flow studies: PerCP-CD45 (Clone: 30-F11 Biolegend), PerCP-Cy5.5- CD3 (Clone: 17A2 Biolegend), ef450-CD4 (Clone: GK1.5 Invitrogen), AF532-Foxp3 (Clone: FJK-16s Invitrogen), Live Dead Aqua (Invitrogen).

### Human HNSCC Single-Cell RNA Sequencing Analysis

Single cell RNA-Seq data (Puram et al., 2017a) was downloaded from the UCSC Cell Browser (Speir et al., 2021) as an expression matrix, metadata, and UMAP coordinates. R software (v4.2.3) was used with package Seurat75 (v4.3.0) to visualize cell populations and import UMAP coordinates and metadata. EphrinB2, SELE, and SELP expression were assessed on cells phenotyped as endothelial cells in the UCSC-provided metadata, and TPM values followed a bimodal distribution where cells could be assumed as ephrinB2 low and high expressors. Log2- transformed fold changes and student’s t-test were calculated to create a volcano plot comparing endothelial cells with high ephrinB2 expression versus low ephrinB2 expression.

### T Cell Isolation and *Ex Vivo* Activation

CD4+ and regulatory T cells were isolated from dissociated spleens using a CD4+CD25+ regulatory T cell isolation kit according to manufacturer’s instructions (Miltenyi Biotec, Bergisch Gladbach, North Rhine-Westphalia, Germany). CD4+ T cells were activated in a 24-well plate (Corning, New York, USA) coated with 2μg/mL anti-CD3 (clone 2C11, Invitrogen, Waltham, Massachusetts) and 2μg/mL anti-CD28 (clone PV-1, BioXCell, Lebanon, New Hampshire, USA). Two days after activation, CD4+ T cells were removed from the coated plate and split to a concentration of 1 x 10^6^ cells/mL. Subsequent splits were performed as needed. 30 U/mL recombinant human IL-2 (Biological Resources Branch National Cancer Institute) was added on day 0 and every other day of activation.

### T Cell Fluorescent Dye-Labeling

7 to 10 days following *ex vivo* activation, T cells were stained with carboxy-fluorescein diacetate succinimidyl ester (CFSE), CellTrace Yellow, or CellTrace Violet proliferation dyes (Invitrogen, Waltham, Massachusetts, USA) at a 1:1000 ratio in serum-free RPMI for 15 to 20 minutes at 37°C. Excess stain was quenched by incubation with 1X volume FBS at room temperature for 2 minutes, then cells were then washed twice with PBS prior to use.

### Intravascular Staining

Staining of lymphocytes in the vasculature was done by intravascular staining as previously described (Estin et al., 2017). APC-conjugated anti-CD4 antibody (clone GK1.5, Biolegend, San Diego, California, USA) was administered by tail vein injection 3 minutes prior to CO2 euthanasia. After euthanizing, perfusions were performed by injection of PBS through the heart to wash out excess stain.

### Extravasation Flow

C57BL/6 and ephrinB2fl/flTie2Cre mice were implanted with 100k MOC2 control or EphB4 shRNA tumors and irradiated 12 to 14 days post-implantation as described above. Two days later, *ex vivo* activated CD4+ T cells were fluorescently stained with CFSE, and 1 x 10^7^ cells were adoptively transferred by tail vein injection. 30 minutes, 60 minutes, 90 minutes, 2 hours, 6 hours, 12 hours, and 24 hours following adoptive transfer, intravascular staining and perfusions were conducted as described above. Tumors were then harvested and processed for flow cytometry as described above. Optimization studies demonstrated that 1 hour was the appropriate time for capturing T cell accumulation in this tumor model. Extravasation flow was therefore performed 1 hour after adoptive transfers for subsequent experiments.

### RNA-sequencing Analysis

Sample preparation and RNA-sequencing analysis were conducted as described previously (Bhatia et al., 2022; Michael W Knitz et al., 2021).

### Boyden Chamber Invasion Assay

MOC2 control or EphB4 shRNA cells were plated to 70% confluency, then the media was replaced with a 1:1 ratio of complete media and Fc-containing conditioned media and the cells were incubated overnight. Cells were then serum starved and 24-well plate inserts with 8µM pores were coated with a 1:10 ratio of Matrigel (Corning) and serum-free media for 4 hours in a 37°C incubator. Following incubation, 750µL complete media containing 10% FBS was added to the bottom chamber, then 500k serum-starved cells were seeded in the Matrigel-coated top chambers. Cells were incubated in a 37°C incubator and allowed to migrate for 24 hours. Next, cells were fixed in 3.7% formaldehyde diluted in PBS for 2 minutes at room temperature followed by 2 washes in PBS. The cells were permeabilized in 100% methanol for 20 minutes, washed twice with PBS, then stained with 0.1% crystal violet for 15 minutes at room temperature. Non-invasive cells on the top of the insert were scraped off with cotton swabs, then the inserts were allowed to dry. Inserts were imaged at 10x magnification, and the number of migrated cells was counted using ImageJ (National Institutes of Health, Bethesda, Maryland, USA). Each condition was performed in five replicates.

### Cancer Cell and CD4+ T Cell Coculture Assay

LY2 control or EphB4 shRNA cancer cells were initially cultured in complete DMEM/F12 medium described above. Subsequently, 100,000 cells were plated overnight in 6-well plates. Cells were then treated with 10 ng/ml of IFN-gamma for 48 hours, after which 10 µg/ml of OVA peptide was added overnight. CD4+ T cells were then isolated from the spleen and lymph nodes of BALB/c mice (DO11.10) using a CD4+ T cell isolation kit according to the manufacturer’s instructions (Miltenyi Biotec, Bergisch Gladbach, North Rhine-Westphalia, Germany). These isolated CD4+ T cells were activated as described above before being co-cultured with LY2 cells at a ratio of 1:1 for 72 hours. Finally, the cells were harvested and stained for flow cytometric analysis.

### Whole-cell lysate preparation

Tumor tissues from indicated group were homogenized in RIPA buffer (Millipore) with a protease inhibitor cocktail (Thermo Fisher Scientific) and phosphatase inhibitors (Sigma-Aldrich) on ice for 30 minutes. Lysates were then collected, and protein concentration was measured using a standard BCA assay as previously described (Bhatia et al., 2016).

### Immunoblot

For western blotting of tumor tissue, proteins were denatured at 95℃ for 7 minutes and then stored at -20℃. Equal amounts of protein (15 μg) were separated using 10% sodium dodecyl sulfate–polyacrylamide (SDS) gels and transferred onto PVDF membranes (#10600023, Amersham Biosciences, Piscataway, New Jersey, USA). To block nonspecific binding sites, the membranes were incubated with 5% Bovine Serum Albumin in Tris-buffered saline containing 1% Tween for 1 hour at room temperature. The membranes were then probed with the specified primary antibodies. Blots were probed overnight at 4°C with respective antibodies. Primary antibodies anti-EFNB2 (#131536, Abcam, Cambridge, United Kingdom), anti-pEFNB2- pTyr316 (SAB 4300631, Sigma), anti-EphB4 (#37-1800, Invitrogen), anti-pEphB4-Tyr987 (PA5- 64792, Invitrogen) and anti-β-actin -HRP conjugate (#5125, Cell Signaling Technology, Danvers, Massachusetts, USA). Horseradish peroxidase (HRP)-conjugated secondary antibodies were obtained from Sigma. Membranes were washed thrice and the antibodies visualized with enhanced chemiluminescent HRP substrate (#R-03031-D25 and R-03025-D25, Advansta, San Jose, California, USA). For detection of signals, X-ray films or versa doc were used. To confirm loading control after detection of phosphorylated protein, membranes were stripped using ReBlot Plus strong antibody stripping solution (#2504, Merck, Germany), blocked and re-probed with different antibodies against whole protein. Protein bands were quantified using *Image lab* & *ImageJ* software. Results are shown as the ratio of total protein or phospho-protein to B actin normalized to the control group. To assign the right protein size, Precision Plus Protein Dual Color Standards Protein Marker was used as a marker (6 μl, #1610374, BIO-RAD).

For immunoblotting of Fc fusion proteins *in* vivo, mouse serum and purified human IgG Fc (#0855911) were diluted in LDS sample buffer (#B0007, Life Technologies, Carlsbad, California, USA) with 2.5% β-mercaptoethanol, heated at 95°C for 2 min, and run on Bolt Bis-Tris Plus gels (#NW04125Box, ThermoFisher Scientific). After semi-dry transfer, the Immobilon membranes were blocked with 5% BSA in 0.1% Tween 20 in Tris buffered saline (TBS) for 1 hour and then incubated in the cold overnight with an anti-human IgG antibody at a 1:1000 dilution (#109-005-098, Jackson ImmunoResearch Labs, West Grove, Pennsylvania, USA). Blots were washed 3 times with 0.1% Tween 20 in TBS before incubating with a horseradish peroxidase-conjugated anti-goat secondary antibody at a 1:4000 dilution (#A16005, Life Technologies) for 1 hour. After 3 washes, the chemiluminescence signal was captured using the ChemiDoc Touch Imaging System (Bio-Rad, Hercules, California, USA). Chemidoc images were quantified using Image Lab (Bio-Rad) to determine the approximate concentration of Fc proteins in the mouse serum using known concentrations of human Fc as standards.

### ELISA

HEK293AD cells stably expressing EphB4 plated on poly-D-Lysine coated plates, or PC3 cells plated without coating, were cultured to 70-80% confluency. Cells were serum starved for 1 hour before treating with ephrinB2-Fc, Fc-TNYL-RAW, or Fc for 30 minutes. Cells were rinsed once with cold DPBS (#21-030-CV, Corning), and collected with lysis buffer (0.5% TX-100 in PBS with Halt Protease and Phosphatase Inhibitor Cocktail (#78443, ThermoFisher Scientific)). After 15 minutes of incubation on ice, cell lysates were centrifuged at 16,000 g at 4°C for 10 minutes, and supernatants were collected. The supernatants were used with the Human Phospho-EphB4 DuoSet IC ELISA kit (#DYC4057-2, R&D Systems, Minneapolis, Minnesota, USA) according to the manufacturer’s instructions.

### Hematoxylin and Eosin Staining and Imaging

Lung samples were submitted to the Dermatology Histology Core at CU Anschutz for cutting and hematoxylin and eosin staining.

### Statistical Analysis

A student’s t-test or one or two-way ANOVA was used when comparing groups for tumor growth curves. A log-rank Mantel-Cox test was used to determine significance in survival studies. The Dunnett post hoc test was used after one-way ANOVA where multiple experimental groups were involved. For western blot analysis, a Mann-Whitney test was performed to compare relative protein expression between control and experimental groups. A Chi-square test was used to when comparing incidence of metastasis between groups. All statistical analysis was done in Prism Software (v9.1.0). Significance was determined by p-values: *p<0.05, **p<0.01, ***p<0.001, ****p<0.0001.

## Acknowledgements

The authors would like to thank the small animal irradiation core at the University of Colorado Anschutz for help designing and implementing the radiation protocols. We would like to thank the University of Colorado Cancer Flow Core Center and the Diabetes Research Center (funded by NIDDK grant #P30-DK116073) for the use of their flow cytometers. We would also like to thank Dr. Jordan Jacobelli and Dr. Rachel Friedman (Barbara Davis Center, University of Colorado Anschutz) for their help in designing a protocol for extravasation flow cytometry. Thank you to Dr. Mohit Kapoor (Schroeder Arthritis Institute, University Health Network, Toronto, ON, Canada) and Dr. Jianping Wu (University of Montreal) for providing us the ephrinB2 knockout mice. Thank you to Dr. Brian Wu and Dr. Philippe Monnier (Krembil Research Institute, University Health Network, Toronto) for their expert advice in knockout mouse generation. Experimental schematics used throughout this work were created with BioRender.com. We would also like to acknowledge our funding sources: R01DE028529, R01CA28465, R01DE028529, 1P50CA261605-01, V Foundation (SDK), F31 DE029997 (LBD), F31 DE033887-01 (JG).

## Supplemental Figure Legends

**Supplemental Figure 1: Loss of EphB4 in cancer cells significantly increases local tumor growth in the context of radiotherapy.** (A) Average tumor volume curves comparing Ctrl shRNA versus EphB4 shRNA MOC2 tumors implanted in C57BL/6 mice. (B) Spaghetti plots of tumor volume for C57BL/6 mice implanted with MOC2 Ctrl (black) or EphB4 shRNA (red) tumors. (C) Dot plot showing significantly larger average tumor volume at 35 DPI in MOC2 implanted C57BL/6 mice. (D) Average tumor volume curves comparing Ctrl shRNA versus EphB4 shRNA LY2 cancer cells implanted in BALB/c mice. (E) Spaghetti plots of tumor volume for BALB/c mice implanted with LY2 Ctrl (black) or EphB4 shRNA (red) tumors. (F) Dot plot showing significantly larger average tumor volume at 20 DPI in LY2 EphB4 shRNA implanted BALB/c mice. (G) Representative 3D lung contouring generated using ITK-Snap from CT scans collected over time for MOC2 implanted C57BL/6 mice. Lung lesions are shown in blue. (H) Hematoxylin and eosin staining of lung tissue for Ctrl and EphB4 shRNA MOC2 implanted C57BL/6 mice. Comparison of tumor volume between the control and experimental group was done using a two-sided student’s *t*-test. Significance was determined if the *p*-value was <0.05*, <0.01**, <0.001***, and <0.0001****. *p*-values are indicated for the figures C \*\*\*\**p* < 0.0001, F \**p* = 0.0002.

**Supplemental Figure 2: Gating strategy for cancer cell and CD4+ T cell coculture experiment.** (A) Gating strategy for flow conducted after coculture of LY2 Ctrl or EphB4 shRNA cancer cells with CD4+ T cells.

**Supplemental Figure 3: Tregs from EphB4 KD tumors downregulate Th1 and Th2 differentiation and mTOR signaling *in vivo*.** (A) Schematic experimental design. Tregs were isolated from spleens and lymph nodes using a MACS Miltenyi Biotec CD4+CD25+ Regulatory T Cell Isolation Kit, and cells were subsequently flow sorted by CD25+ expression. B) Bulk proteomic pathway analyses showing GO processes that were upregulated in Tregs of MOC2 EphB4 shRNA compared to Ctrl tumor implanted C57BL/6 mice. (C) Bulk proteomic pathway analyses showing KEGG pathways that were downregulated in Tregs of MOC2 EphB4 shRNA compared to Ctrl tumors implanted in C57BL/6 mice. (D) Bulk proteomic pathway analyses showing GO processes that were downregulated in Tregs of MOC2 EphB4 shRNA compared to Ctrl tumor implanted C57BL/6 mice.

**Supplemental Figure 4: EphrinB2 KO in vascular endothelial cells coupled with radiation therapy reduces local tumor growth.** (A) Spaghetti plots comparing tumor volume between WT hosts and ephrinB2^fl/fl^Tie2^Cre^ mice implanted with MOC2 WT tumors.

**Supplemental Figure 5: EphrinB2 expression in vascular endothelial cells increases after RT.** (A) RNA expression of ephrinB2 in sorted VE-Cadherin+ vascular endothelial cells 6 hours after 5 Gy total body irradiation of C57BL/6 mice. (B) Quantification of ephrinB2-expressing vascular endothelial cells (CD45-CD31+PDPN-) in the TME of MOC2 tumors treated with or without radiation. (C) Quantification of Ki-67 expression in ephrinB2-expressing vascular endothelial cells in the MOC2 TME. 10 Gy was administered 7 DPI, and tissue was harvested 17 DPI. Comparison of differences between the control and experimental group was done using a two-sided student’s *t*-test. Significance was determined if the *p*- value was <0.05*, <0.01**, <0.001***, and <0.0001****. *p*-values are indicated for the figures B \**p* = 0.0373, C \**p* = 0.0285.

**Supplemental Figure 6: Gating strategy for immune cell populations in the TME of ephrinB2 KO mice.** (A) Flow cytometry gating strategy for tumor immune cell populations in WT hosts or ephrinB2^fl/fl^Tie2^Cre^ mice implanted with 100k MOC2 WT cancer cells in the buccal mucosa.

**Supplemental Figure 7: EphrinB2 KO in vascular endothelial cells affects the systemic immune response.** (A) Flow cytometry gating strategy for the blood compartment. (B) Quantification of granzyme B expressing CD8+ T cells in the blood. (C) Flow cytometry gating strategy for the DLN compartment. (C) Quantification of CD4+ T cells in the DLNs. (E) Quantification of CD8+ T cells in the DLNs. (F) Quantification of dendritic cells in the DLNs. (G) Quantification of Tregs in the DLNs. Comparison of differences between the control and experimental group was done using a two-sided student’s *t*-test. Significance was determined if the *p*-value was <0.05*, <0.01**, <0.001***, and <0.0001****.

**Supplemental Figure 8: Specialized flow cytometry demonstrates increased CD4 T cell accumulation in the TME of ephrinB2 KO mice.** (A) Schematic of flow cytometry extravasation experiments designed to examine CD4+ T cell accumulation into the TME. EphrinB2^fl/fl^Tie2^Cre^ mice or WT controls were implanted with 100k MOC2 EphB4 shRNA cells. 14 days post-implantation, mice were irradiated with 10 Gy. 2 days later, 10 million CD4 T cells were stained and adoptively transferred. At various timepoints following adoptive transfers, tail vein injections of CD4-APC were administered to label vascular immune cells. After 3 minutes, mice were euthanized, and their hearts were perfused to flush out excess stain. Tumors were subsequently harvested for flow cytometric analysis. (B) Quantification of intravascular and intratumoral CD4+ T cells at various timepoints following adoptive transfers. Graph on the left shows a close-up view of shorter timepoints where tumors were harvested 30, 60, and 90 minutes following adoptive transfer. (C) Quantification and ratios of intravascular and intratumoral CD4 T cells 1 hour following adoptive transfer into ephrinB2^fl/fl^Tie2^Cre^ or WT hosts implanted with EphB4 shRNA tumors. Comparison of differences between the control and experimental group was done using a two-sided student’s *t*-test. Significance was determined if the *p*-value was <0.05*, <0.01**, <0.001***, and <0.0001****. *p*-values are indicated for the figures C \**p* = 0.0141.

**Supplemental Figure 9: EFNB2-Fc-His and Fc-TNYL-RAW-GS reduce local tumor growth.** (A) Spaghetti plots comparing tumor volume among C57BL/6 mice implanted with MOC2 WT cancer cells, treated with 8 Gy RT, and transfected with plasmids by hydrodynamic tail vein injections. Mice treated with RT alone (black) or SB + RT (green) served as controls. Experimental mice were treated with EFNB2-Fc-His (blue) or Fc-TNYL-RAW-GS (purple).

**Supplemental Figure 10: Detection of Fc fusion proteins in mouse serum and EphB4 phosphorylation following treatment with EFNB2-Fc-His and Fc-TNYL-RAW-GS.** (A) Blood was collected from tumor-bearing mice 26 days after plasmid hydrodynamic tail vein injection and the indicated nanoliters (nl) of serum were separated by SDS-PAGE and probed by immunoblotting with anti-human Fc antibodies. The indicated amounts of purified human Fc were also included for comparison and to enable approximate quantification of the Fc fusion proteins from mouse blood. Serum was obtained from 3 mice injected with each construct. (B) ELISA measuring EphB4 tyrosine phosphorylation induced by 2.5 nM purified commercial ephrinB2-Fc (ephrinB2-Fc R&D), EFNB2-Fc-His, Fc-TNYL-RAW-GS peptide, or Fc purified from the culture medium of transfected HEK293 cells. Phosphorylation was measured for endogenous EphB4 expressed in PC3 prostate cancer cells (left) or EphB4 stably expressed in HEK293 cells (right). Fc was used as a control.

