## supplemental figures for "Manipulating the EphB4-ephrinB2 axis to reduce metastasis in HNSCC"

### Supplemental Figure 1: Loss of EphB4 in cancer cells significantly increases local tumor growth in the context of radiotherapy.

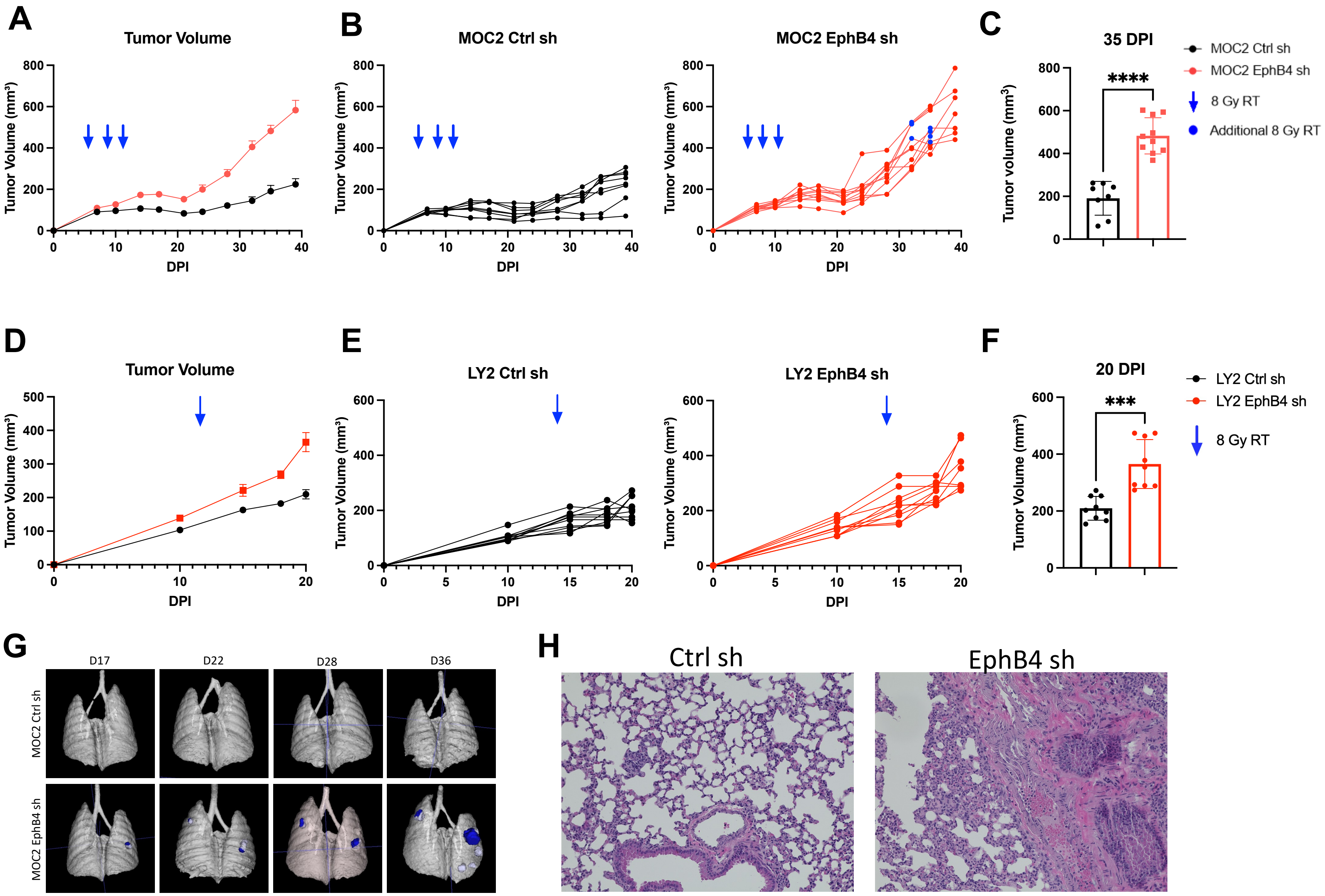

#### Supplemental Figure 2 : Gating strategy for cancer cell and CD4+ T cell coculture experiment

**A**

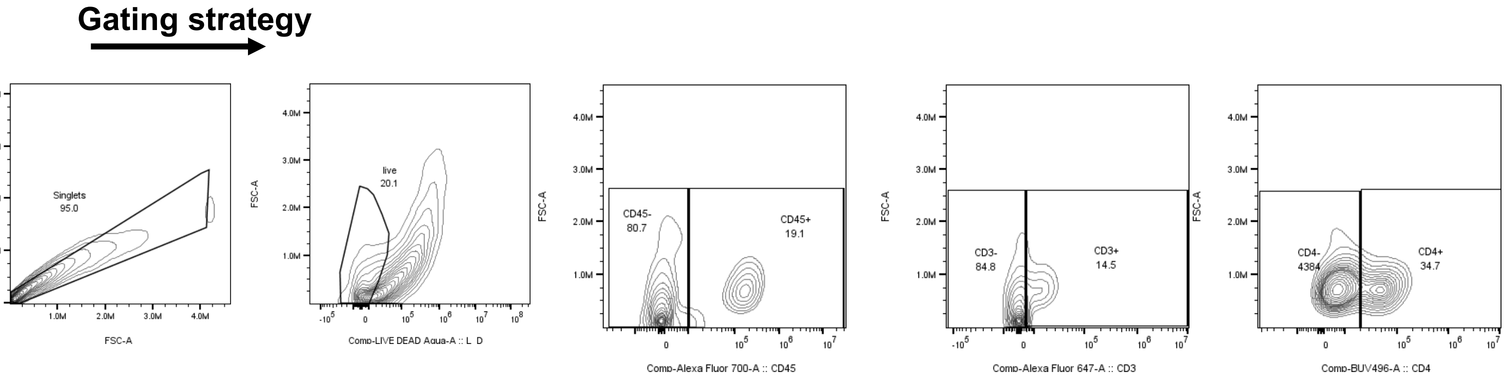

**Supplemental Figure 3: Tregs from EphB4 KD tumors downregulate Th1 and Th2 differentiation and mTOR signaling *in vivo*.**

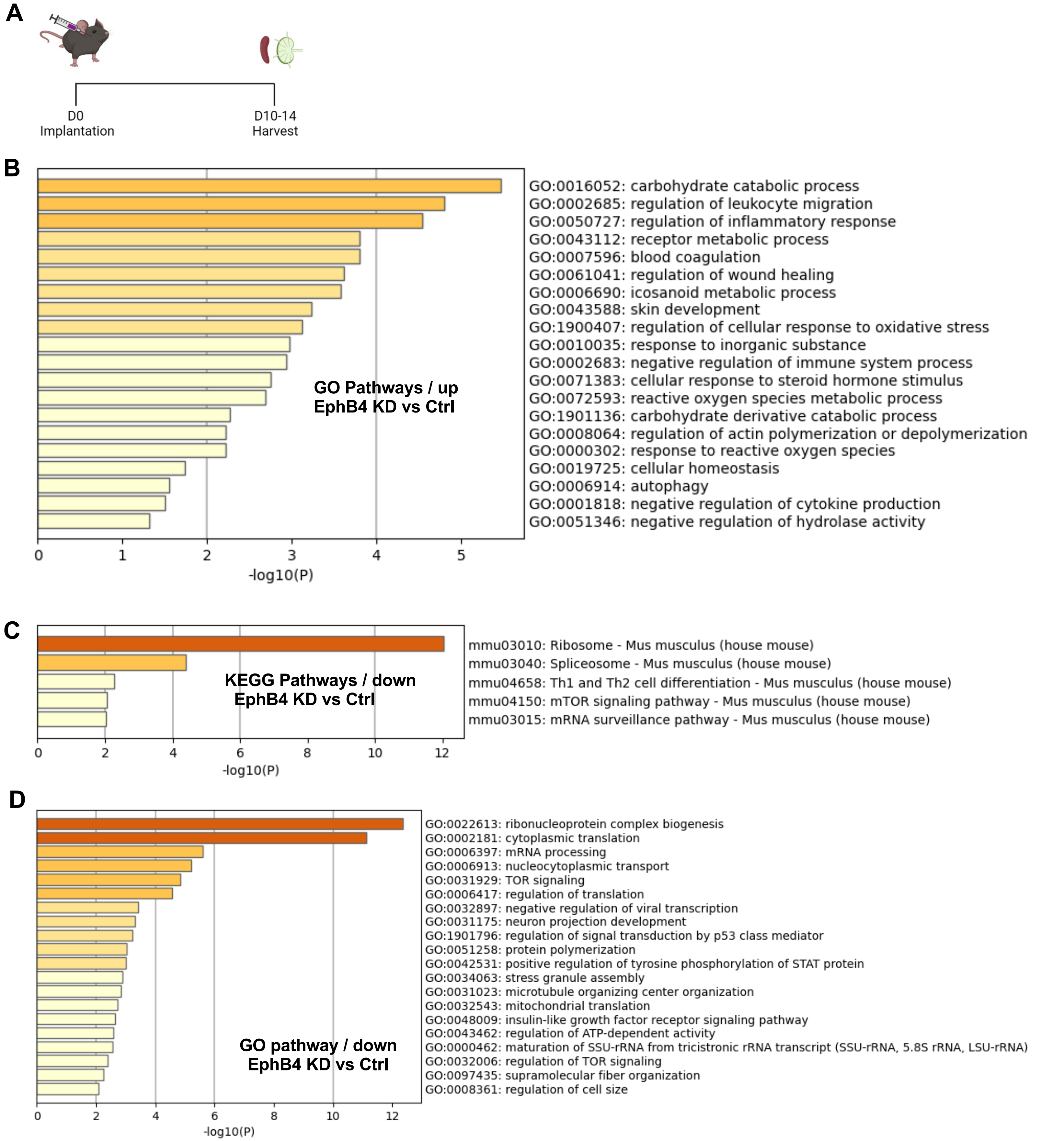

**Supplemental Figure 4: EphrinB2 KO in vascular endothelial cells coupled with radiation therapy reduces local tumor growth.**

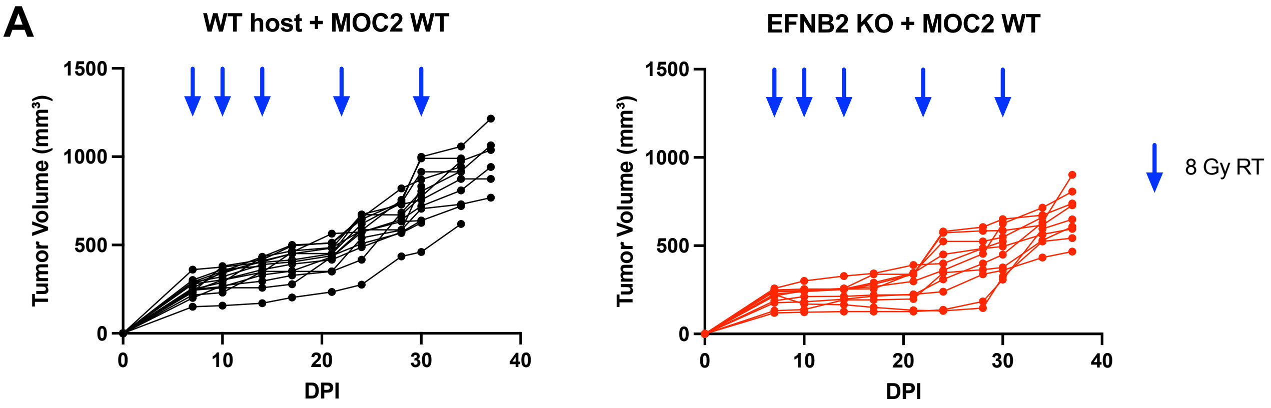

**Supplemental Figure 5: EphrinB2 expression in vascular endothelial cells increases after RT.**

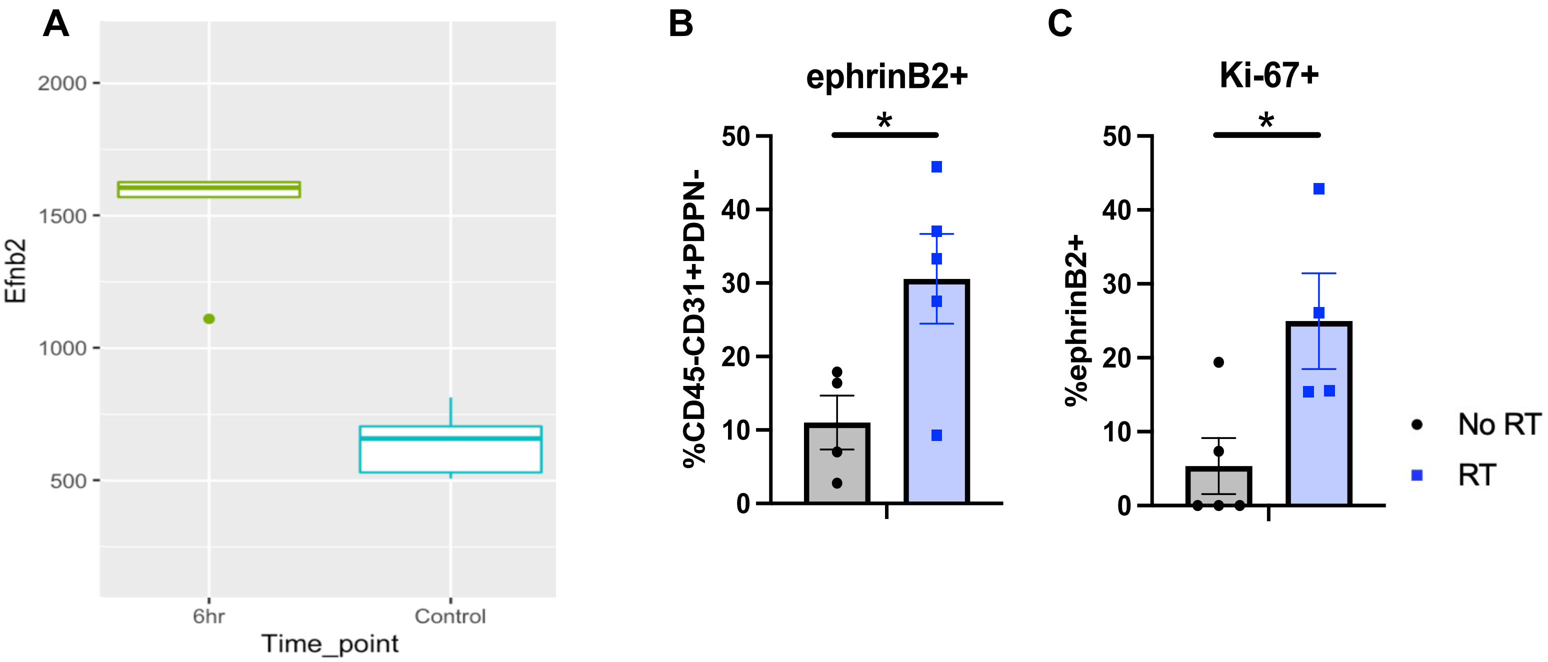

**Supplemental Figure 6: Gating strategy for immune cell populations in the TME of ephrinB2 KO mice.**

**A**

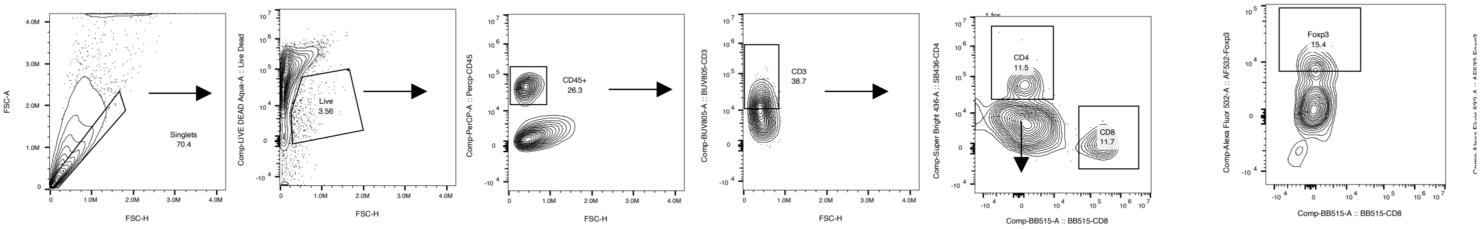

### Supplemental Figure 7: EphrinB2 KO in vascular endothelial cells affects the systemic immune response.

**A** **Blood gating**

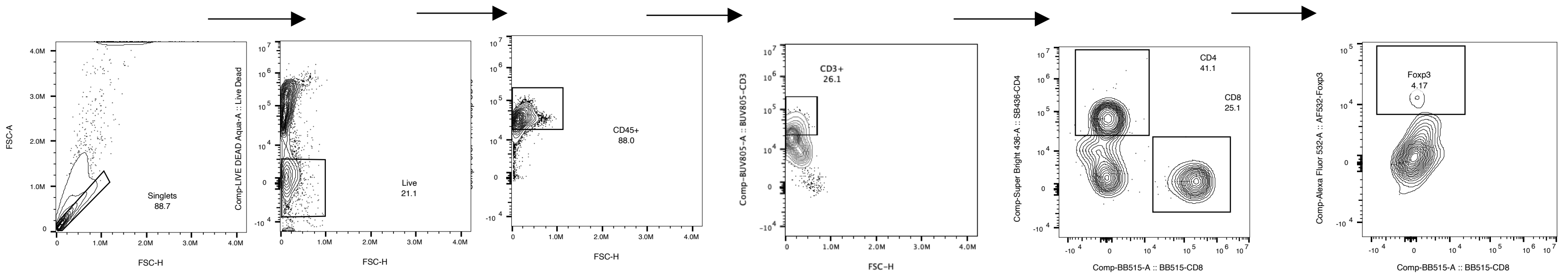

**B**

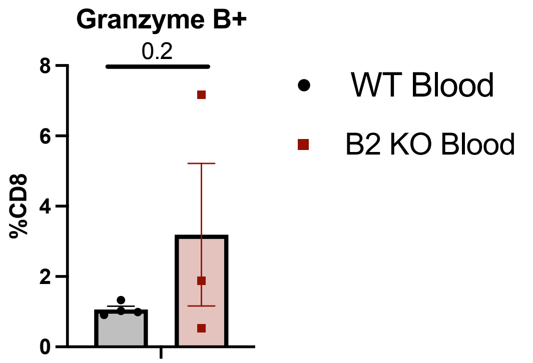


**C** **DLN gating**

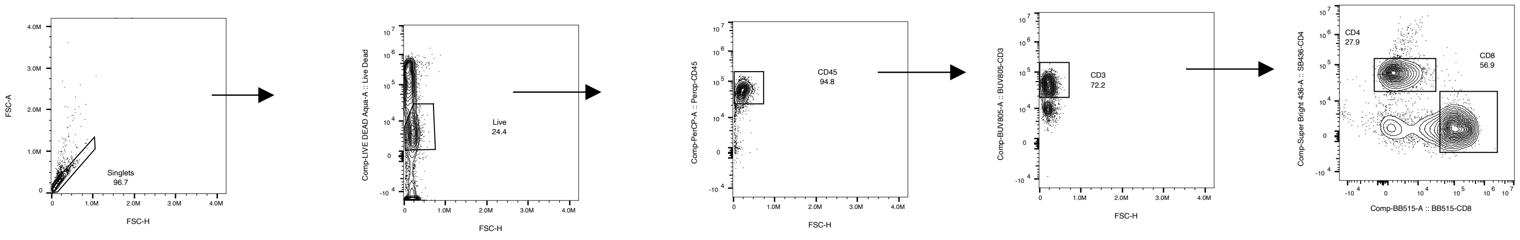

**D**

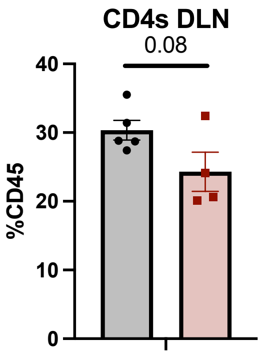


**E**

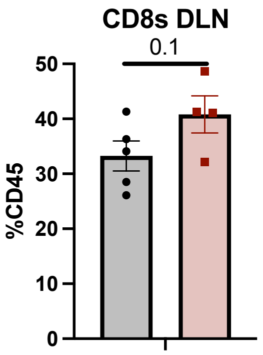


**F**

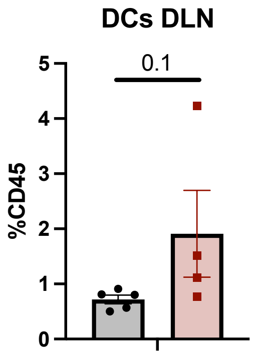


**G**

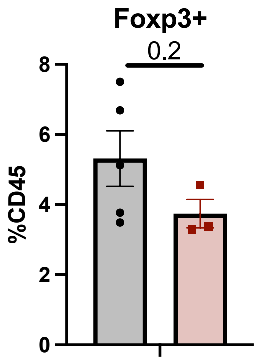


WT DLN

B2 KO DLN

**Supplemental Figure 8: Specialized flow cytometry demonstrates increased CD4 T cell accumulation in the TME of ephrinB2 KO mice.**

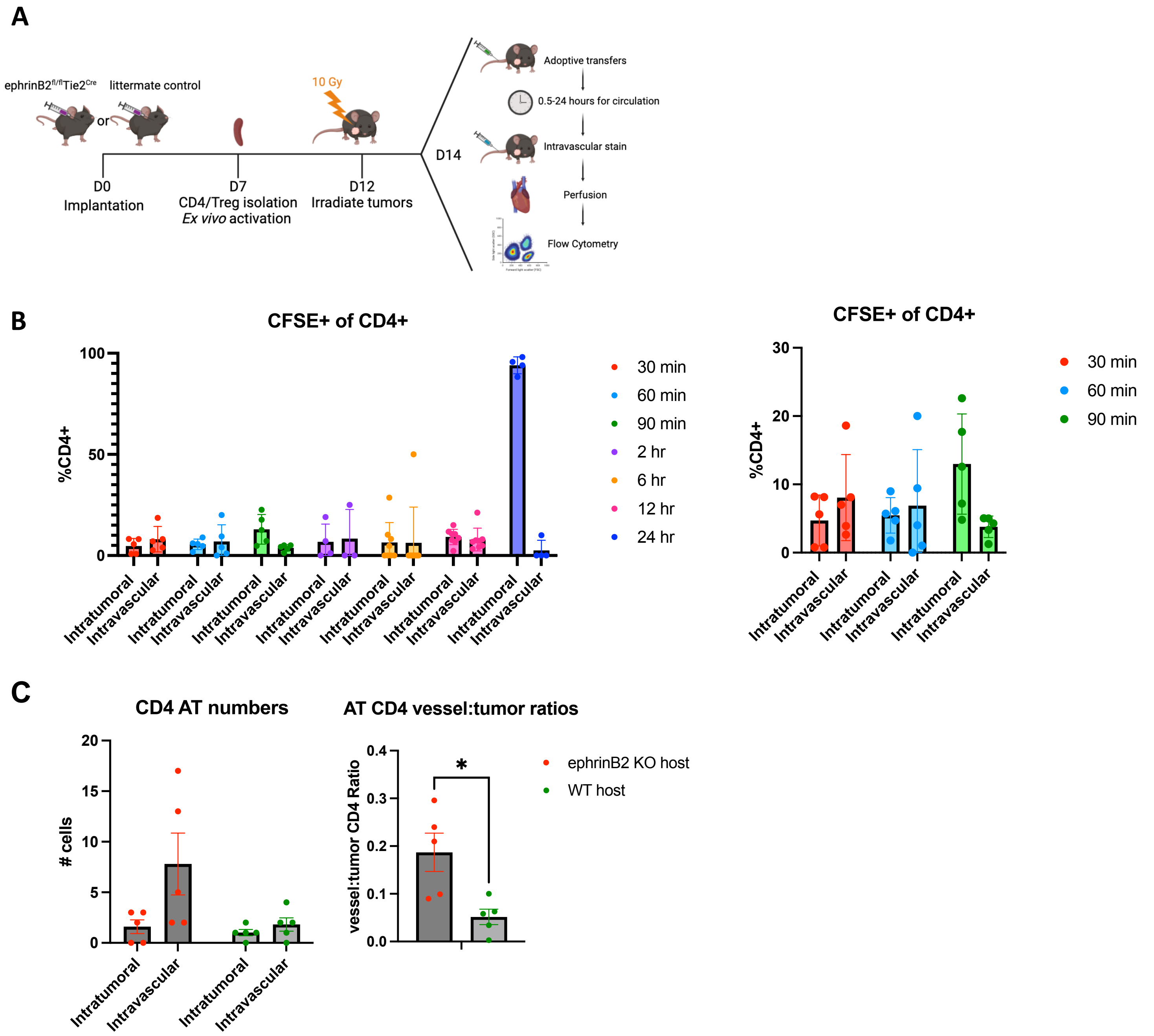

### Supplemental Figure 9: EFNB2-Fc-His and Fc-TNYL-RAW-GS reduce local tumor growth.

**A**

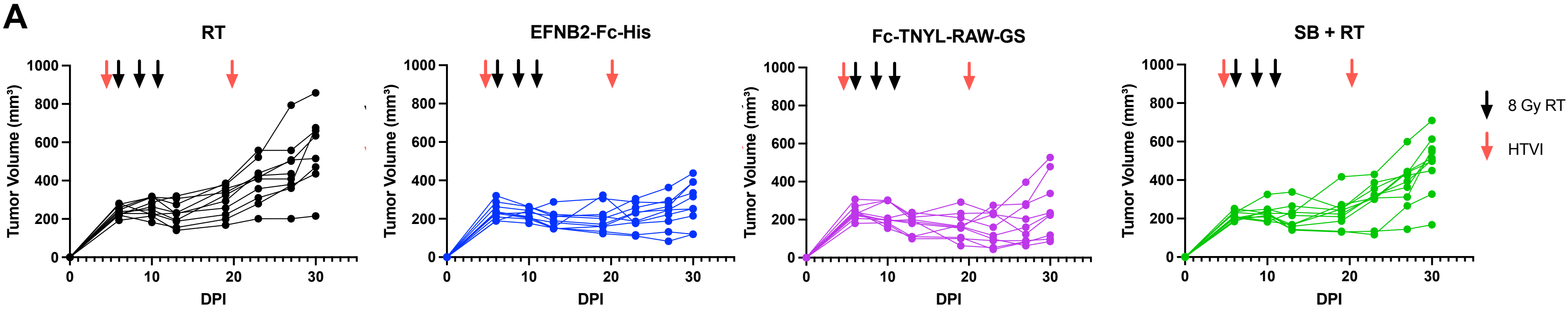

**Supplemental Figure 10: Detection of Fc fusion proteins in mouse serum and EphB4 phosphorylation following treatment with EFNB2-Fc-His and Fc-TNYL-RAW-GS.**

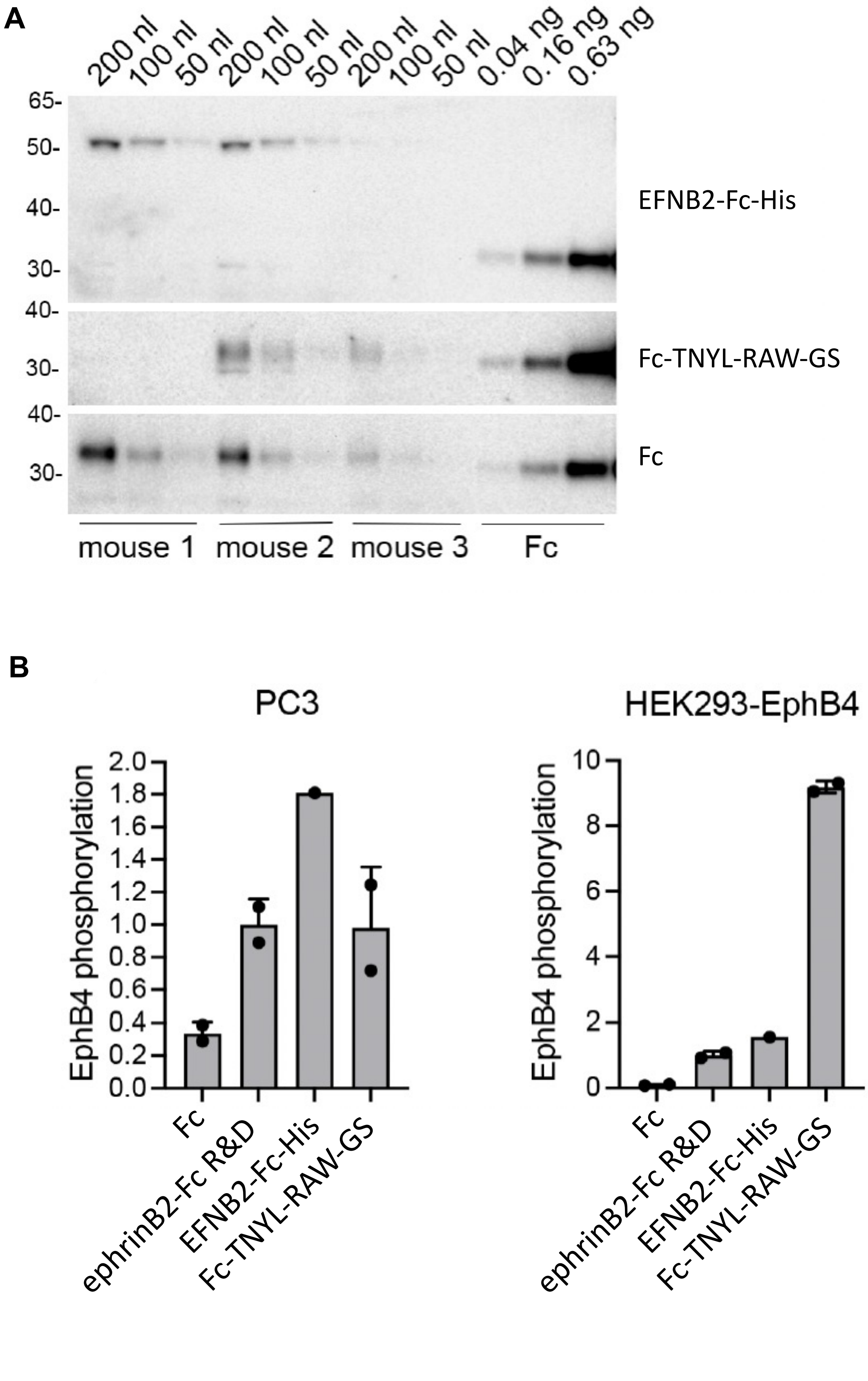
